# Single-Cell RNA-seq Uncovers Dynamic Processes Orchestrated by RNA-Binding Protein DDX43 in Chromatin Remodeling during Spermiogenesis

**DOI:** 10.1101/2022.06.12.495783

**Authors:** Huanhuan Tan, Weixu Wang, Chongjin Zhou, Yanfeng Wang, Shu Zhang, Pinglan Yang, Rui Guo, Wei Chen, Lan Ye, Yiqiang Cui, Ting Ni, Ke Zheng

**Affiliations:** State Key Laboratory of Reproductive Medicine, Nanjing Medical University, Nanjing 211166, China; State Key Laboratory of Genetic Engineering, Collaborative Innovation Center of Genetics and Development, Human Phenome Institute, Shanghai Engineering Research Center of Industrial Microorganisms, School of Life Sciences and Huashan Hospital, Fudan University, Shanghai 200438, China

## Abstract

Advances in single-cell RNA sequencing (scRNA-seq) have allowed for elucidating biological mechanisms at cell state level. Mammalian spermatogenic process showcases dynamic switches of gene expression pattern with delicate morphological and functional alterations of germ cells, but it is unclear how such dynamics is genetically controlled. Here we demonstrate that mouse testis-enriched RNA helicase DDX43, as well as its ATP hydrolysis site, is required for spermiogenesis. Genetic mutation of *Ddx43* renders spermatids heterogeneously defective in multiple steps of chromatin remodeling, resulting in incomplete substitution of transition protein by protamine and less condensed sperm nucleus. Through scRNA-seq analyses of testicular cells derived from adult wild-type and *Ddx43* mutant testes in mice, we reveal that the DDX43 deficiency-elicited perturbation in the dynamic RNA regulatory processes underlies the differentiation deficiency of spermatids. Further, focused analyses on early-stage spermatids combined with enhanced CLIP sequencing (eCLIP-seq) identify *Elfn2* as DDX43-targeted hub gene, whose *in vivo* knockdown shows similar phenotypic defects as *Ddx43* mutant. Our study illustrates an essential role for DDX43 in post-meiotic chromatin remodeling and highlights the single cell-based strategy for a refined dissection of stage-specific regulation of germline differentiation.

## Introduction

After meiosis, haploid spermatids undergo spermiogenesis, which is characterized by a cascade of chromatin remodeling events that culminate in the replacement of most histones by protamines^1–5^. This step-wise process entails massive histone modifications as well as transient formation of DNA strand breaks (DSBs)^6–9^. Genomic DNA is compacted to high orders, giving rise to a firm sperm head with greatly condensed nucleus that ensures stable intergenerational transmission^6, 10^. While causality is well-established linking histone-to-protamine transition to male fertility^4, 11, 12^, recent studies begin to give mechanistic insights into how this transition is specifically and directly controlled by several important proteins^6, 7, 13–17^. It’s conceivable that many more genes may coordinate this long multi-step developmental journey. Particularly, it remains unclear whether there’s an RNA regulator that governs transcriptomic dynamics to orchestrate chromatin remodeling.

Round spermatids differentiate through eight typical steps till they become elongating and then elongated, during which transcription is being converted gradually from active to inert^18–20^. Previous studies often selected postnatal time points to enrich or even purify certain types of germ cells for either dynamics characterization^21–24^, or for an inter-genotypic comparison^25–28^. As total RNA-seq used in these studies only reveal an ensemble average level of a mixed cell population at a fixed time point, such traditional strategies pose a difficulty in disentangling profound cellular heterogeneity. To put it differently, the static snapshots of the morphology-based, but not intrinsic, molecular alterations do not allow for tracing the footprint of contiguous cell states. In recent years, single-cell RNA-seq (scRNA-seq) has provided an effective avenue for discriminating a vast number of individual germ cells based on their differential transcriptomic patterns^29–34^. For example, scRNA-seq has depicted the functional distinction between round spermatid subpopulations in their potential for embryo development, reflecting intrinsic subtypes^29^. To achieve developmental lineage tracing of single cells, some computational tools, such as ICAnet^35^ and scVelo^36^, are built effective for the inference of differentiation hierarchy and trajectory. By virtue of single cell-based methodologies, it’s therefore promising to elucidate how inter-subtype transitions of varying transcriptomic states of spermatids are regulated, presumably by an RNA-binding protein^37, 38^.

RNA helicases are RNA-binding proteins with versatile functionality^39, 40^. While the majority of RNA helicases are expressed across multiple tissues, some play specific roles in the germline^41–44^. We notice that DEAD-box helicase protein DDX43 is highly expressed in testis and many tumors^45, 46^. Whereas *in vitro* biochemical studies have demonstrated DDX43 serves as a *bona fide* helicase^47^, its function in the testis remains unknown. In the present study, we integrated multiple approaches to query the biological and molecular roles of DDX43 in spermatogenesis. Genetic mutation of *Ddx43* leads to deficiency in sperm chromatin compaction and disturbs a cascade of chromatin remodeling events. Using scVelo to infer germ cell developmental trajectory and velocity-directed Markov chain simulation to predict the differentiation terminal, we found that spermatids have a pronounced alteration in the differentiation trajectory, in concert with their histochemical defects, in *Ddx43* mutant mice. Coupling with eCLIP-seq, we anchored an early subpopulation of post-meiotic spermatids as initial targets of DDX43 and further identified *Elfn2* as a hub gene downstream of DDX43. These findings together reveal DDX43 operates an early gene expression network to maintain the homeostasis of subsequent chromatin dynamics in spermatids.

## Results

### DDX43 expression dominates late spermatocytes and early spermatids

To begin characterizing DDX43 distribution, western blot analysis of mouse tissues showed that DDX43 was not ubiquitously expressed, but testis-enriched (Fig. 1a), consistent with the results detected by quantitative reverse transcription followed by polymerase chain reaction (qRT-PCR) (Extended Data Fig.1a). In the testis, *Ddx43* transcript levels increased sharply at postnatal day 14 (P14) (Extended Data Fig.1b). In parallel, DDX43 protein abundance increased sharply at P21, a time point when late-stage spermatocytes differentiate into round spermatids, and maintained at higher levels thereafter (Fig. 1b). Specifically, it was abundant predominantly in pachytene spermatocytes and secondly in round spermatids (Fig. 1c and Extended Data Fig.1c), shown from both protein and mRNA levels in these two types of isolated germ cells (Extended Data Fig. 1d, e, f). Immunofluorescence in adult testis sections, as well as western blot in subcellular fractions, revealed that DDX43 protein distributed over nucleus and cytoplasm (Fig. 1d and Extended Data Fig.1g). Co-immunofluorescence of Lectin-PNA, an acrosomes marker, allowed us to visualize DDX43 spatiotemporally dynamic from stage I to XII of seminiferous tubules and accordingly across different steps of germ cells, e.g., step 1-16 spermatids (Fig. 1d). In detail, DDX43 was predominantly cytoplasmic and slightly nuclear in early spermatocytes (stage I-IV), whereas its nuclear portion progressively increased in late spermatocytes, was robust at meiotic metaphase cells (stage XII), and then decreased in post-meiotic round spermatids (Fig. 1d). DDX43 was weakly detected in preleptotene (stage VII-VIII), zygotene (stage XI-XII), and at much higher levels in pachytene (stage I-IV), continuously rising until the expression peaked at meiotic metaphase cells, and was still present in early round spermatids, but decreased after step 8 (stage VIII) (Fig. 1d). The immunofluorescence specificity of DDX43 was proved by comparing with mice homozygous for *Ddx43* knockout allele (*Ddx43*^−/−^; see details in next paragraph) as negative control (Fig. 1e). Taken together, DDX43 is nucleocytoplasmic and mostly expressed before and after meiotic exit (Fig. 1f), implicating its major function around this timing.

**Figure 1.**
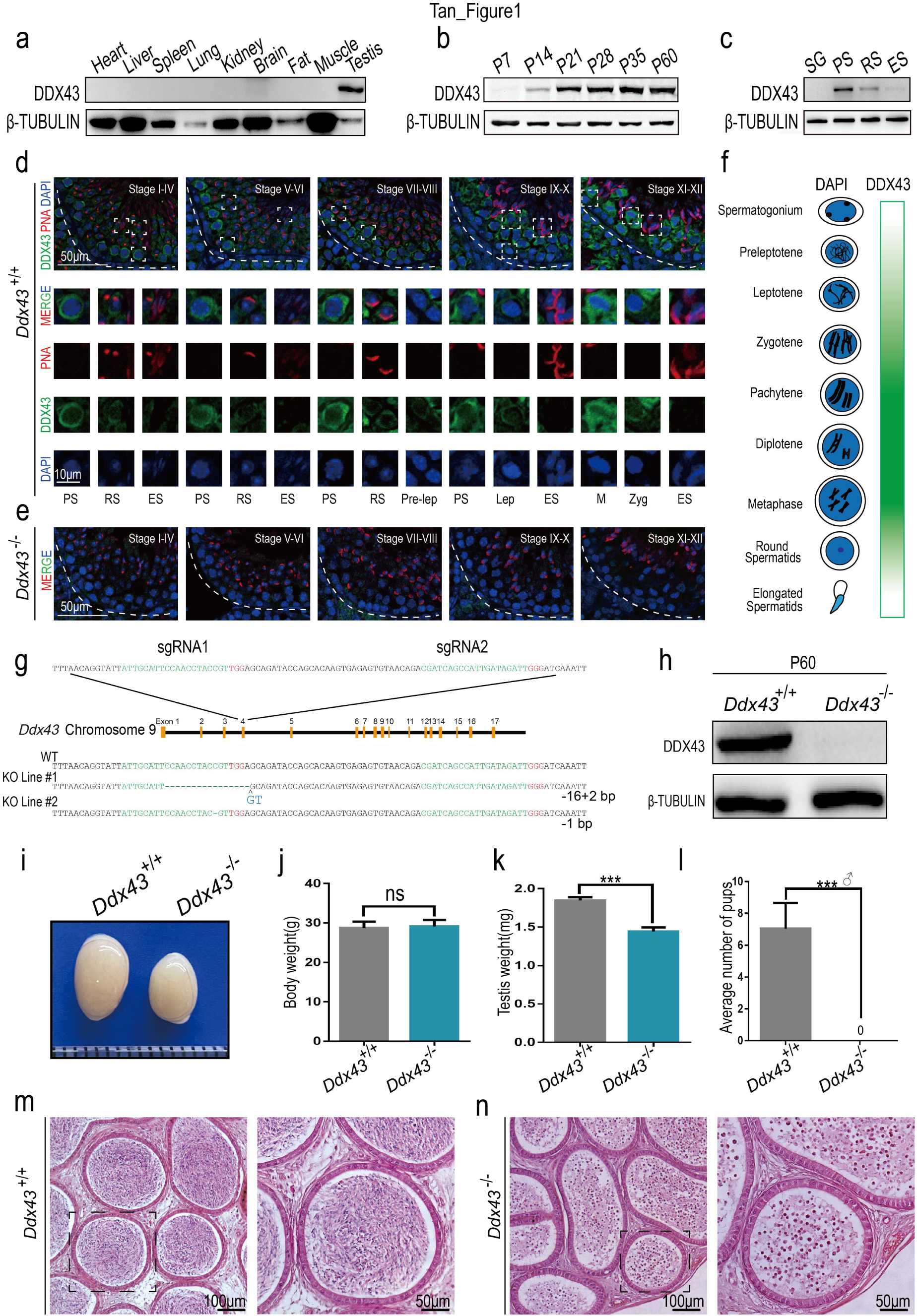
DDX43 is enriched in mouse testis and indispensable for male fertility. (a-c) Western blot analyses of DDX43 protein in extracts from P60 mouse tissues (A), testes collected from mice at different ages (B), and purified spermatogenic cells (C), including spermatogonia (SG), pachytene spermatocytes (PS), round spermatids (RS), and elongating spermatids (ES). β-TUBULIN serves as a loading control. (d, e) Immunofluorescence staining of DDX43 (green), PNA (red), and DAPI (blue) on sections from adult wild-type (d) and *Ddx43^−/−^* (e). Lower panels show magnifications of the boxed area in the upper panels. PS, pachytene; RS, round spermatids; ES, elongating spermatids; Pre-lep, pre-leptotene; Lep, leptotene; Zyg, zygotene; M, metaphase cells. Scale bars indicated are applicable for all panels within a group. (f) Graphic representation of developmental expression pattern of DDX43 in male germ cells. (g) Schematic diagram illustrating the CRISPR-Cas9 genome editing strategy. PAM sequences are in red. sgRNA targeting sites are in green. Mutant sequence are in blue. (h) Western blot analyses confirm absence of DDX43 protein in P60 *Ddx43^−/−^* mice testes lysates. β-TUBULIN serves as an internal loading control. (i) Representative images from P60 mice testes indicating atrophied testes in the *Ddx43^−/−^* mice. (j-l) Comparative analyses of body weight (j), testes weight (k), and fertility (l) between adult *Ddx43^+/+^* and *Ddx43^−/−^* mice. Male mice were naturally crossed with age-matched wild-type female mice. n=5 for each genotype, ***P < 0.001; ns, not significant (Student’s t-test). (m, m) Hematoxylin and Eosin (H&E) staining of epididymis sections from adult *Ddx43^+/+^* (m) and *Ddx43^−/−^* (n) mice. Scale bar is indicated.

### Spermiogenesis is defective in *Ddx43* mutant mice

To elucidate the biological function of *Ddx43*, we first generated two *Ddx43* knockout (KO) lines by CRISPR-Cas9 gene editing approach. Two small guide RNAs (sgRNAs) were used to target exon 4 of *Ddx43* gene (Fig. 1g), producing two mutant founder lines (Fig. 1g and Extended Data Fig. 2a). KO Line #1 has an insertion of 2-bp (base pair) accompanied by a 16-bp deletion; KO Line #2 has a deletion of a single bp. Both alleles result in reading frameshifts and premature termination codons. Genotyping PCR and RT-PCR confirmed the absence of full-length transcripts in mice homozygous for the mutant allele (hereafter referred to as *Ddx43*^−/−^), indicating successful targeting of *Ddx43* gene (Extended Data Fig. 2b, c, d). Western blot and immunostaining analyses demonstrated absence of DDX43 protein in testes from adult *Ddx43*^−/−^ mice (Fig. 1e, h). Mice of all *Ddx43* genotypes were matured to adulthood normally without apparent developmental defects. Compared with age matched wild-type mice, the body weight was similar (Fig. 1j), but *Ddx43*^−/−^ testis looked smaller with a lower weight (Fig. 1i, k). Histologically, in contrast to wide-type epididymis teemed with spermatozoa (Fig. 1m), germ cell numbers in *Ddx43^−/−^* mice were drastically reduced and no normal spermatozoa were present; instead, sloughed irregular spermatids and cell debris were observed (Fig. 1n). Consistently, no pups were produced from adult *Ddx43*^−/−^ males in mating tests (Fig. 1l).

Considering DDX43 is an explicit ATP-driven RNA helicase in possession of RNA binding and unwinding activities^47–49^, and a potential mediator of dynamic protein-RNA interactions^40^. That means knockin of a point mutation (DQAD) within DEAD-box will inactivate DDX43 ATP hydrolysis and lock its bound RNA targets from their turnover. Hence, an *in vivo* fascinating mechanism is worth probing for DDX43 serving as an RNA helicase, like the case of MVH^50, 51^. To this end, we generated a *Ddx43* knockin model (referred to as *Ddx43*^KI^) (Fig. 2a), and obtained the KO Line #3 by chance. Adult *Ddx43*^KI/KI^ mice phenocopied *Ddx43^−/−^* mice in male fertility, manifested with smaller testis (Fig. 2d), male sterility (Fig. 2e), no normal mature sperm (Fig. 2b), yet, comparable body weight (Fig. 2c). While the RNA abundance of *Ddx43* in knockin testes were still comparable to wild type (Extended Data Fig. 3a, b), it is regrettable that DDX43 protein was undetectable not only in total lysates (Extended Data Fig. 3c), unless after immunoprecipitation (IP) enrichment (Extended Data Fig. 3d), but also in cellular subfractions (Extended Data Fig. 3e, f). The fact that the catalytic-dead mutant protein, but not mutant RNA, was dramatically reduced in *Ddx43*^KI/KI^ testis reflects ATP hydrolysis-dependent self-regulation of the DDX43 protein. While the helicase-specific role of DDX43 is masked by the unexpected ablation of the full catalytic-dead protein, we still could disclare that mutation of the DDX43 ATP hydrolysis site result in DDX43 deficiency and a failure of normal sperm production. For this reason, we focused on these two models for subsequent studies.

**Figure 2.**
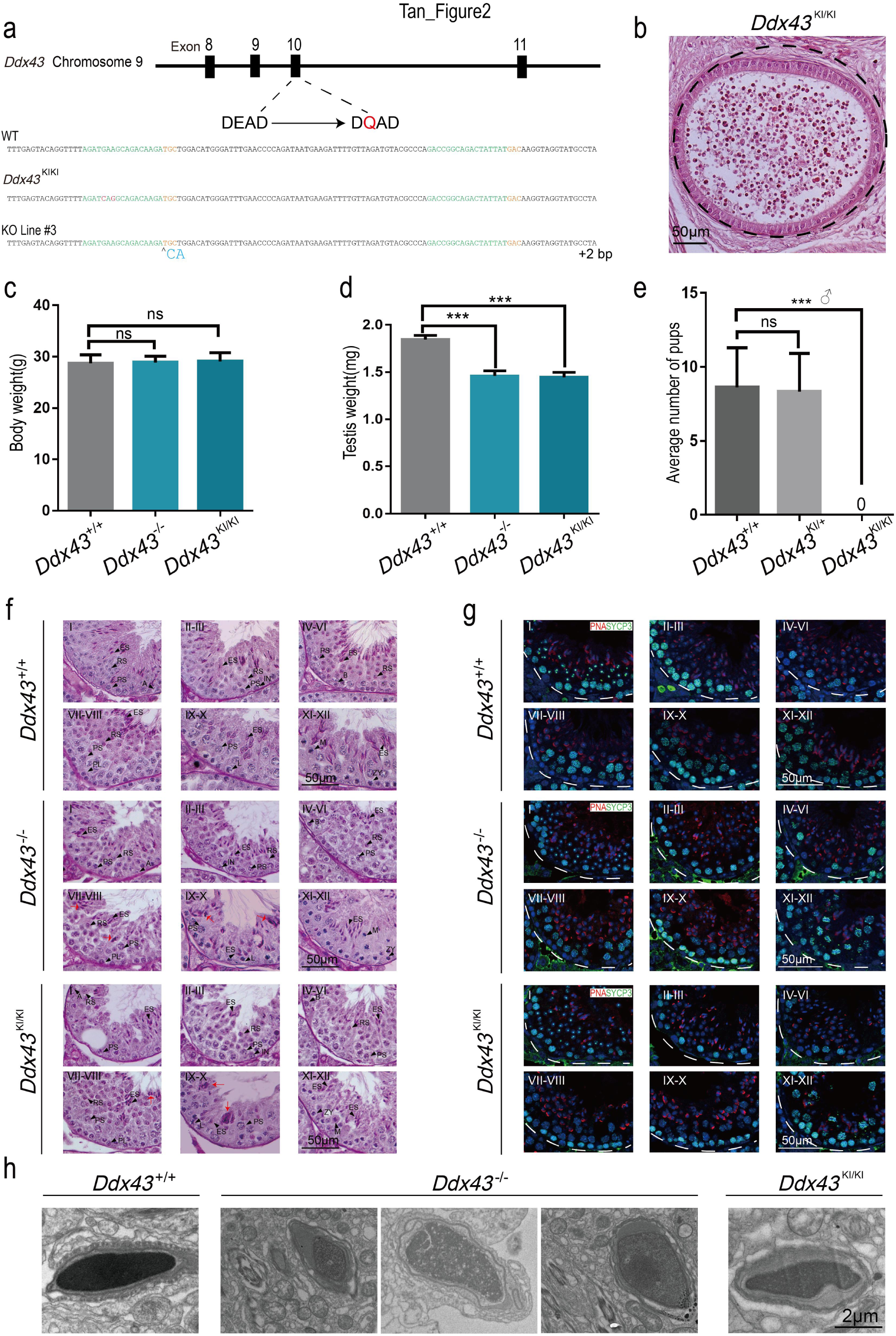
DDX43 deficiency impairs chromatin compaction during spermiogenesis. (a) Using CRISPR-Cas9 to create the catalytic-dead *Ddx43* mice that carry a point mutation in the ATPase motif from DEAD to DQAD. (b) H&E staining of epididymis sections from adult *Ddx43*^KI/KI^ mice. Both *Ddx43^−/−^* and *Ddx43*^KI/KI^ epididymis are void of normal spermatozoa. Scale bar is indicate (c-d) Comparative analyses of body weight (c), testes weight (d) between adult *Ddx43^+/+^* and *Ddx43*^KI/KI^ mice. (e) Fertility test result for adult *Ddx43^+/+^*, *Ddx43^KI/+^* and *Ddx43*^KI/KI^ mice naturally crossed with age-matched wild-type female mice. n=5 for each genotype, ***P < 0.001; ns, not significant (Student’s t-test). (f) PAS-hematoxylin staining of testes sections from adult *Ddx43^+/+^*, *Ddx43*^KI/KI^ and *Ddx43^−/−^* mice, Stages of seminiferous tubule are denoted by Roman numerals. A, type A spermatogonia; B, type B spermatogonia; ES, elongating spermatids; IN, intermediate spermatogonia; L, leptotene spermatocytes; M, metaphase cells; PS, pachytene spermatocytes; PL, preleptotene spermatocytes; ZY, zygotene spermatocytes. Red arrows represent abnormal spermatids at step 16. Scale bar is indicated. (g) Immunofluorescence staining of SYCP3 (green) and PNA (red) for further staging of spermatogenesis. DNA was counterstained with DAPI. Scale bar is indicated. (h) Representative images of transmission electron microscopy of sperm heads from adult mice. Scale bar is indicated.

We performed histological examination with testes sections and noticed germ cells in mutant testes displayed comparable behavior at P21, differentiation differences at P28, a time point coinciding with the appearance of elongating spermatids, and pronounced flaws at P35 (Extended Data Fig. 2e). To define defects in each step of germ cells, we performed staining using Periodical-Aid-Schiff (PAS) (Fig. 2f) and co-immunostaining for SYCP3 and PNA in adult testis sections (Fig. 2g). *Ddx43*^−/−^ and *Ddx43*^KI/KI^ seminiferous tubules presented fewer elongated spermatids relative to wild type, and lacked normal spermatozoa (Fig. 2f, g). At stage VIII, most *Ddx43* mutant step 16 spermatids showed a failure of normal spermiation with irregular head shape and excess cytoplasm (Fig. 2f; red arrows). These condensed rod- or round-like abnormal elongated spermatids frequently appeared in stages IX-XI (Fig. 2f; red arrows). In *Ddx43* mutant epididymis, although there were only an extremely low number of sperm, we still enriched a few and observed their nuclei were more efficiently stained with acidic aniline (Extended Data Fig. 2f), indicative of less condensed chromatin in these mutant sperm. Transmission electron microscopy (TEM) further revealed that sperm heads in *Ddx43* mutant mice were less condensed with uneven density and malformed structure (Fig. 2h). Taking account all these observations, we conclude that DDX43 deficiency leads to defects in spermiogenesis and chromatin compaction in sperm.

### DDX43 deficiency disturbs a cascade of molecular events in chromatin remodeling

During spermiogenesis, transition proteins (TNPs) replace canonical histones and are subsequently substituted by protamines (PRMs)^3, 15^. This switch produces a greater than six-fold condensation of somatic chromosomes, resulting in a packaged chromatin structure^2, 10^. To address the potential cause whereby genetic mutation of *Ddx43* compromises chromatin compaction during spermiogenesis, we embarked on inspection of chromatin remodeling events by co-immunolabeling TNPs and PNA. At stage XII-I, aside from the difference in the morphology of some elongating spermatids in adult *Ddx43*^+/+^, *Ddx43*^−/−^ and *Ddx43*^KI/KI^ testes, TNPs in testes of all genotypes displayed similar fluorescent localization and intensity on their nuclear heads (Extended Data Fig. 4a, b). At stage VIII, whereas TNPs were normally missing in wild-type step 16 elongated spermatids, both transition protein 1 (TNP1) and 2 (TNP2) were present in partial *Ddx43^−/−^* and *Ddx43*^KI/KI^ seminiferous lumens (Fig. 3a, b). And, the majority of TNP-positive cells presented irregular rod- or round-like heads (Fig. 3a, b; white arrows), in contrast to TNP-negative cells (Fig. 3a, b; gray arrows). At the same stage VIII, although protamine 2 (PRM2) in some *Ddx43* mutant elongated spermatids was as intensively visible as wide type, it was often absent or fragmentary on many others, not as uniformly visible as in nearly all wide-type elongated spermatids (Fig. 3c). More importantly, some condensed spermatids were retained at stage IX, when the TNP2 signals on the less condensed chromatin were conspicuous (Fig. 3d; white arrows) and those on the more condensed were weak or absent (Fig. 3d; gray arrows). In parallel, at stage X, the retained spermatids were mainly accompanied with positive signals (Fig. 3e, white arrows), and partially negative (Fig. 3e; gray arrows). These special phenomena at stage IX-X match well with our aforementioned shot in Fig. 2f. In addition, TNP2 retention was found in the sporadic premature sperm sloughed to *Ddx43*^−/−^ epididymis (Extended Data Fig. 4c). Taken together, these data indicate that DDX43 deficiency impairs TNP-to-PRM substitution, although this transition is not absolutely blocked.

**Figure 3.**
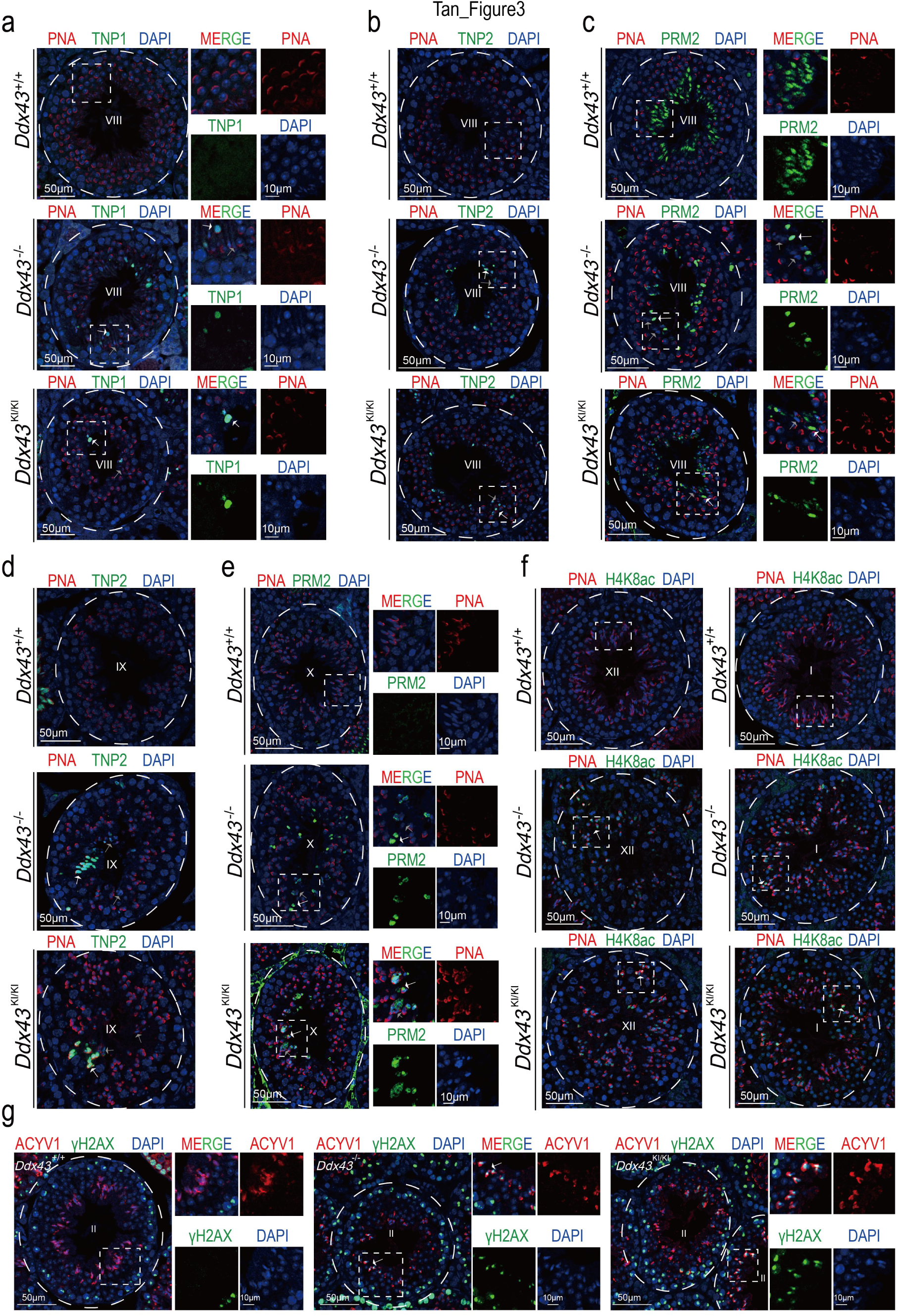
DDX43 deficiency impairs chromatin remodeling during spermiogenesis. (a-g) Immunofluorescence analyses in adult *Ddx43^+/+^*, *Ddx43*^KI/KI^ and *Ddx43^−/−^* mouse testes with the following combinations. (a) TNP1 (green) and PNA (red), See also Extended Data Fig 4a; (b, d) TNP2 (green) and PNA (red), See also Extended Data Fig 4b; (c, e) PRM2 (green) and PNA (red); (f) H4K8ac (green) and PNA (red), See also Extended Data Fig 5a, b; (g) γH2AX (green) and PNA (red). Right panels show magnifications of the boxed area in the left panels (a, c, e, g). Stages of seminiferous tubule are denoted by Roman numerals. White arrows indicate positive signals; gray arrows indicate negative signals. DNA was counterstained with DAPI. For all images above, stage numbers and scale bars are indicated.

Hyperacetylation of histone H4 takes place just prior to TNP replacement of canonical histones and is thought to be a crucial step for the subsequent events^19, 52, 53^. In accordance with previous report^6, 52^, in wide-type testes, a strong labeling for H4K8ac was confined to step 9-11 elongated spermatids at stage IX-XI (Fig. 3f and Extended Data Fig. 5a). In *Ddx43* mutant testes, while signal intensity of H4K8ac was comparable at stages IX-X (Extended Data Fig. 5b), H4K8ac-positive cells remained there at stages XII-I (Fig. 3f; white arrows). The partial eviction of H4K8ac in mutant late elongating spermatids prompted us to further examine the phosphorylated H2AX (γH2AX), a marker for transient physiological DNA strand breaks (DSBs), which is pivotal for the process of nucleosomal DNA supercoils elimination and DNA damage response^8^. In wild-type testis, γH2AX was cleared in elongated spermatids from step 13 (stage I) onward (Fig. 3g). In *Ddx43* mutant testes, γH2AX was still present in step 14 (stage II) elongated spermatids (Fig. 3g; white arrows). In order to directly assess DSBs, the sensitive TUNEL assay was conducted^9, 54, 55^. As a result, there was a significant increase in TUNEL-positive signals at step 13-16 in *Ddx43^−/−^* compared with wild-type testis (Extended Data Fig. 5b), implying the retention of γH2AX is likely attributed to uncleared DSBs. These results suggest DDX43 is required for hyperacetylated H4 eviction and DNA repair during spermiogenesis.

### DDX43 deficiency is insufficient to frustrate meiosis and pachytene piRNA biogenesis

Considering the onset of DDX43 expression from spermatocytes as well as the potential defects in DSB repair in *Ddx43* mutant spermatids, DDX43 deficiency might as well impact meiotic events such as DNA double-strand breaks (DDSBs) and chromosome synapsis^56, 57^. To this end, we conducted co-immunofluorescence analysis of SYCP3 (a lateral element of the SC complex^58^) and γH2AX on the chromosome spreads of spermatocyte nuclei. To our expectation, γH2AX staining patterns are indistinguishable for each type of spermatocyte between wide-type and mutant testes (Extended Data Fig. 6a). Nor obvious defects were observed via stage-specific comparison of double immunofluorescence for SYCP1 (a central element of the SC complex^59^) and SYCP3 (Extended Data Fig. 6b), combined patterns of which are well-adopted to define each stage of the meiotic prophase I. In concert with the robust production of post-meiotic round spermatids, these results suggest that DDX43 has no major influence on meiotic progression.

DDX43 was recently reported to function as a Vasa-like protein to facilitate amplification of Piwi-interacting RNAs (piRNAs) in an ovary-derived cell line of silkworms^60^. In mammalian testis, pachytene piRNAs, as one predominant class of piRNAs, are generated in pachytene spermatocytes and round spermatids^61, 62^. Based on its expression pattern, DDX43 might regulate pachytene piRNA biogenesis in mouse testis. To this end, we radiolabeled total small RNAs and nevertheless showed comparable abundance of pachytene piRNAs in wild-type versus mutant testis (Extended Data Fig. 7a). As expected, immunofluorescence did not show derepression of two major piRNA-associated retrotransposons, long interspersed nuclear element-1 (LINE1, L1) retrotransposon-derived open reading frame 1 (ORF1) and intracisternal A-particle (IAP) in mutant testes (Extended Data Fig. 7b). MIWI and MILI, two classical piRNA pathway proteins, normally localize to the inter-mitochondrial cement (IMC) of spermatocytes and concentrated in chromatoid bodies (CBs) of round spermatids^63, 64^. As expected, neither their protein levels (Extended Data Fig. 7c) nor localization patterns (Extended Data Fig. 7d) were affected. Thus, unlike in silkworms, DDX43 seems dispensable for piRNA biogenesis in mice.

### RNA velocity captures differentiation deficiency events induced by *Ddx43* mutation

Next, we performed RNA-seq using wild-type and *Ddx43* mutant testes (three per genotype) collected at P21 and P28, time points separately enriched for spermatocytes and spermatids. Counting up those expressed in either genotype, there were total 14,972 and 15812 transcripts at P21 and P28 in *Ddx43^−/−^*, as well as 14965 and 15856 in *Ddx43^KI/KI^*, respectively. Among them, by contrast, only 21 and 109 expressed genes in *Ddx43^−/−^* testes, as well as 34 and 115 in *Ddx43^KI/KI^* testes, were significantly altered (P < 0.05, fold change > 2) (Extended Data Fig. 6c). In view of the nature of DDX43 being an RNA helicase, the limited number of differentially expressed genes above raised a puzzle for us to seek what is indeed responsible for DDX43-mediated chromatin remodeling during spermiogenesis. We speculate that because total RNA-seq data reflect rather an ensemble average of different cell types, it may limit a deep deciphering of the highly dynamical regulation in the process of chromatin remodeling.

To explore the impact of DDX43 on gene expression at cell type resolution, we performed 10x Genomics single-cell RNA-seq (scRNA-seq) throughout the complete spermatogenic lineage in 12580 testicular cells derived from adult wild-type, DEAD-box point mutation knock-in (*Ddx43^KI/KI^*) and *Ddx43* knock-out (*Ddx43^−/−^*) testes (two per genotype). To assess both the cell typic compositional changes and transcriptional regulation differences, we need to build a high confidential transcriptome reference manifold based on our wild-type samples, and then project the *Ddx43^KI/KI^* and *Ddx43^−/−^* samples on this reference. After doing quality control (Method), using our recently developed single cell clustering method independent component analysis based gene co-expression network inference (ICAnet)^35^, combined with previously described cell-type specific markers^34^, we identified all major germ cell populations covering whole development process, including spermatogonia (SPG), meiotic spermatocytes (SCytes), post-meiotic haploid round spermatids (STids), and elongating spermatids (ES) (Fig. 4a, b). We also detected three somatic cell compartments includes Endothelial, Macrophage and an Unknown somatic cell type (Fig. 4a, b). To support our cell type identifications, we compared our results with previous fluorescence activated cell sorting (FACS) based smart-seq2 scRNA-seq data ^29^, and the correlation heatmap supported the accuracy of our annotations (Fig. 4c). We also noticed that SCytes2 act as the transition state between SCytes and STids. To further support our annotation, we quantified the metagene expression of differential expression gene sets between MII and the early stage of STids (up-regulated genes in MII and up-regulated genes in RS1o2) in the cells of SCytes2 and STids using AUCell^65^. As expected, in general, the MII metagene have higher expression value in SCytes2 (Extended Data Fig. 8a). On this basis, we further calculated each cell type’s differentiation entropy^66^, which is regarded as one of the best characterization metrics for cell differentiation potency, and noticed that the differential entropy showed perfectly gradient descent along the differentiation trajectory (Fig. 4d; SPG -> SCytes -> STids -> ES). Collectively, above in-depth computational analyses supported the accuracy of our cell type annotation and differentiation trajectory, which means we have an ideal latent space to capture the majority transcriptional regulation information during the spermatogenesis so that we could address the perturbation effects posed by *Ddx43* mutation at cell type and/or state resolution.

**Figure 4.**
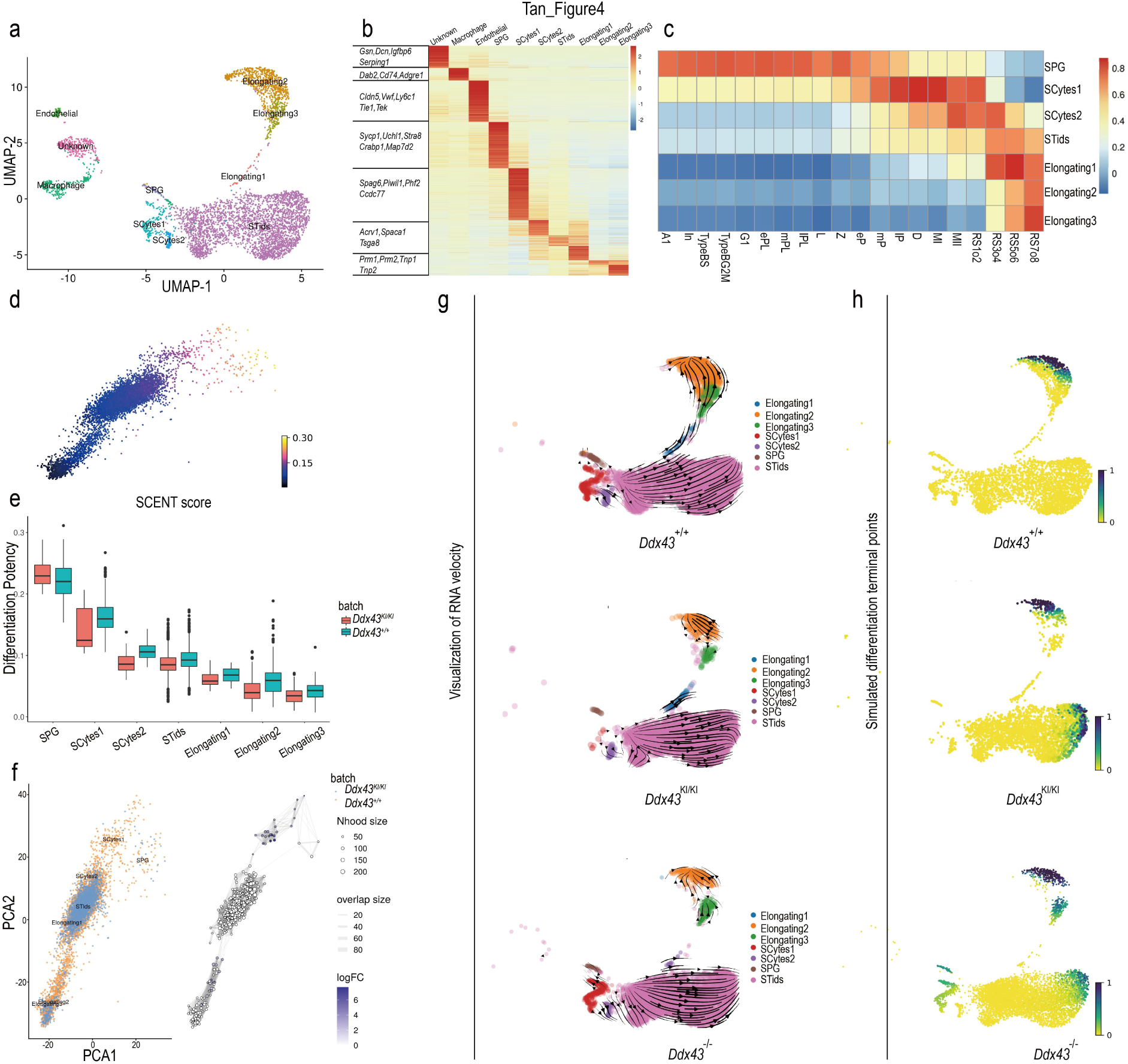
Overview of Single-cell transcriptome profiling on adult *Ddx43^+/+^* and *Ddx43* mutant whole testes. (a) UMAP and clustering analysis of combined single-cell transcriptome from mice testes (n = 9250). Each dot represents a single cell and is colored according to its cluster identity as indicated on the figure key. The 10 cluster identities were assigned based on marker gene expression. UMAP: Uniform Manifold Approximation and Projection. SPG, spermatogonia; SCytes, meiotic spermatocytes; STids, post-meiotic haploid round spermatids; ES, elongating spermatids. (b) The heatmap for expression of selected marker genes for the 10 cell types. (c) The heatmap of cell type gene expression spearman correlation between our annotated single cell reference expression profile with flow-sorted single cell gene expression profile. A1, type A1 spermatogonia; In, intermediate spermatogonia; BS, S phase type B spermatogonia; BG2, G2/M phase type B spermatogonia; G1, G1 phase preleptotene; ePL, early S phase preleptotene; mPL, middle S phase preleptotene; lPL, late S phase preleptotene; L, leptotene; Z, zygotene; eP, early pachytene; mP, middle pachytene; lP, late pachytene; D, diplotene; MI, metaphase I; MII, metaphase II; RS2, steps 1–2 spermatids; RS4, steps 3–4 spermatids; RS6, steps 5–6 spermatids; RS8, steps 7–8 spermatids. (d) PCA plot of single cell transcriptome data with cells colored based on their cell differentiation potency (network entropy) calculated by SCENT. (e) The boxplot of cell differentiation potency for each differentiation stages during spermatogenesis of both *Ddx43^+/+^* and *Ddx43*^KI/KI^ mice (f) A neighborhood graph of the results from Milo differential abundance testing (right panel). Nodes are neighborhoods, colored by their log fold change across genotypes. Non-differential abundance neighborhoods (P-value > 0.05) are colored white, and sizes correspond to the number of cells in each neighborhood. Graph edges depict the number of cells shared between neighborhoods. The layout of nodes is determined by the position of the neighborhood index cell in the PCA (left panel). (g, h) Visualization of the RNA velocity analysis results on the UMAP plot (g, left panel), and predicted terminal points based on velocity directed trajectory (h, right panel). Cells are colored according to their differentiation terminal probability.

After building the wild-type reference of mouse spermatogenesis process, we mapped the query cells from *Ddx43^KI/KI^* and *Ddx43^−/−^* sample on the reference by an anchor-based integration methods^67^, and transferred the reference label to each query cell (Method). Both marker gene expression pattern and flow-sorted experimental data supported the accuracy of our annotation on query cells (Extended Data Fig. 8b, d). We also applied the metagene analysis in query cells to support the correct annotation of transition cell type SCytes2 (Extended Data Fig. 8c, e). One of the critical tasks of scRNA-seq is to shed light on the abnormal trajectory events in perturbated sample, which makes the cell states deviate from normal. We calculated the differentiation potency of mapped mutant cells, and noticed that there exists clearly reduced differentiation capacity starting from the stage of SCytes to the ES in *Ddx43^KI/KI^* sample (Fig. 4e) and *Ddx43^−/−^* sample(Extended Data Fig. 8f). This deficiency was identified through evaluating expression pattern of each cell, and may also reflect on the manifolds as the population density of cell type changes during the conversion of each type. In order to visualize the population-level dynamics, we next used a novel differential abundance testing method^68^ to visualize the cell population density perturbation on the sampled local region of cell k-nearest neighbored (kNN) graph, and we identified differential abundant neighborhoods from the stage SCytes2 to stage STids, and stage STids to ES (Fig. 4f, and Extended Data Fig. 8g), indicating that STids was the crucial cell type/stage relevant to the abnormal cell differentiation in *Ddx43* mutant testes.

A further question is whether *Ddx43* deficiency changes the differentiation trajectory of spermatogenic cells. As splicing is a continuous process, a spliced transcript at a given time point is likely derived from an unspliced transcript at a previous time point. We therefore applied scVelo, a model evaluates RNA splicing velocity by fitting the chemical master equation function through Expectation-Maximization (EM) algorithm to derive a highly dimensional vector predictive of future transcriptional state of individual cells^36^, to infer developmental trajectory of spermatogenesis based on single-cell RNA-seq data^69^. Notably, *Ddx43* deficiency changed the direction of the velocity vector within the STids. Unlike wild-type, *Ddx43* mutant STids showed weak progression towards ES (pointed by most arrows) (Fig. 4g). To make a clearer visualization of the perturbation effects, we built a Markov chain and used velocity vector information as the orientation of transition matrix, so that we could predict the terminal differentiation points on the manifold that is reflected by a terminal probability of each cell. Interestingly, in wild-type testes, the cells could safely differentiate to the final stage, ES, while in *Ddx43^KI/KI^* and *Ddx43^−/−^* testes, some cells were trapped in the stage of STids (Fig. 4h), in line with our immunohistochemistry observation (Fig. 2f, g). Taken together, these analyses imply that *Ddx43* deficiency misled the differentiation path of STids.

### DDX43 operates dynamic RNA regulatory processes at early-stage STids

Encouraged from the above analyses, we zoomed in the STids for re-clustering and found four sub-clusters (Fig. 5a). To gain insights into the gene expression dynamics along these four clusters, we analyzed the sub-cluster specific expression genes (Fig. 5b and Extended Data Fig. 9a). Specifically, genes expressed at higher levels in sub-cluster 0 are enriched in RNA processing, DNA repair, chromosome segregation and chromatin remodeling. Genes enriched in ribosome biogenesis are at higher levels in sub-cluster 1. Sub-cluster 2 is enriched for genes involved in proteasomal protein modification and processing, spermatid differentiation and acrosome reaction. And, sub-cluster 3 is enriched for genes that control fertilization, sperm-egg recognition, sperm motility and flagellated sperm motility. These gene ontology terms were in accordance with previous studies about the serial differentiation in mouse male germ cells^29, 34^. To connect those sub-clusters, we ordered cells in a pseudo-temporal manner using PAGA^70^ and RNA velocity^36^, and noticed the differentiation direction starts from sub-cluster 0 towards sub-cluster 2 (Fig. 5c, d).

**Figure 5.**
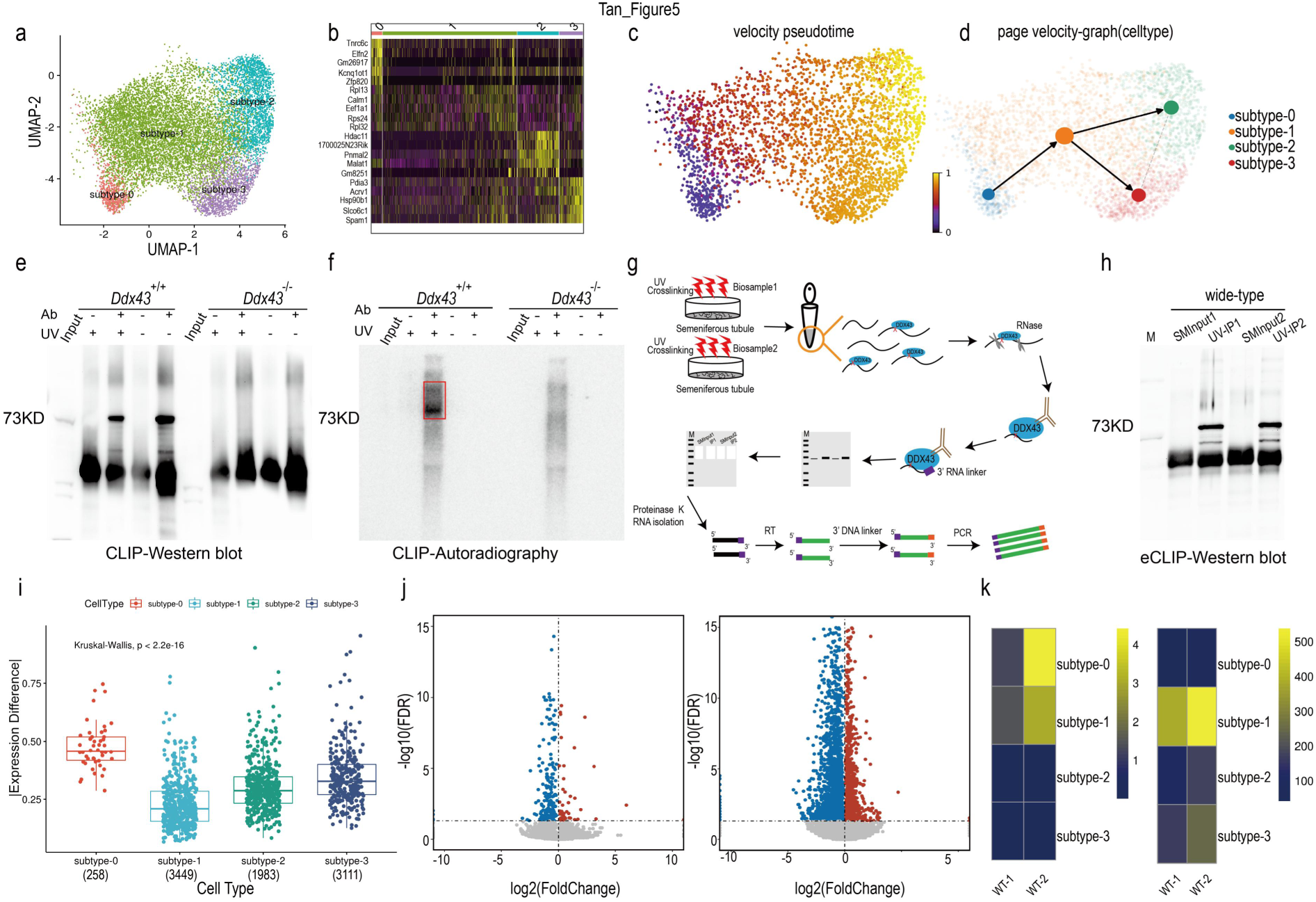
Identification of four transcriptional states for STids with scRNA-seq and eCLIP-seq. (a) Zoom-in analysis of STids reveals four cellular states (subtype0 to subtype3) during spermatid development. (b) Top 5 highly expressed genes for each subtype in STids. (c) Velocity Pseudotime analysis on STids cells. Each cell is colored according to its velocity time. (d) PAGE directed graph of four states. (e) Western blot analyses of DDX43-RNA complexes prepared by P^32^-radioactive conventional CLIP in P24 testes. Both non-crosslinked and *Ddx43^−/−^* samples served as negative controls. (f) Autoradiography of the conventional CLIP complexes from panel e. (g) Schematic of enhanced CLIP (eCLIP) protocol. See the Methods section for details. (h) Western blot validation of DDX43-RNA complexes prepared by eCLIP in P24 wild-type testes. SMInput1 and SMInput2 serve as negative controls in two biological replicates, respectively. (i) The boxplot for absolute expression difference value of *Ddx43* differential expression targets (DETs) in four wide-type cellular states. (j) Volcano plot showing gene differential expression of subtype0 and subtype1. (k) The heatmap of DDX43 targets enrichments in down-regulated genes of two eCLIP-seq replicates. The heatmap is colored according to the -log10 (P-value). We used Fisher-exact test to calculate statistical significance of enrichments (left panel). The heatmap of the number of DETs in two eCLIP-seq replicates (right panel).

Then, we asked which sub-cluster may play a crucial role in regulating the differentiation process of STids. To answer this question, we attempted to detect RNA directly bound by DDX43 *in vivo*. We radiolabeled DDX43-bound RNA after UV-crosslinking immunoprecipitation (CLIP) in P24 testis lysates. Western blot analysis and autoradiography clearly showed CLIP enrichment of DDX43 protein and in parallel its size-matched RNA in wild-type, but not in *Ddx43^−/−^* testes (Fig. 5e, f), supporting that our antibody could efficiently capture crosslinked DDX43-RNA ribonucleoproteins after stringent washes. To achieve a robust profiling of DDX43-bound RNAs at a transcriptome-wide level, we applied enhanced CLIP (eCLIP) as before ^71, 72^(Fig. 5g). Western blot analysis confirmed eCLIP enrichment of DDX43 protein (Fig. 5h). ‘Size-matched input’ (SMInput) served as control libraries to reduce nonspecific background from eCLIP libraries. We processed DDX43 eCLIP reads and successfully mapped them using an established pipeline ^71^(Extended Data Fig. 10a), exhibiting high correlation between biological and technical replicates (Extended Data Fig. 10b). High reproducibility of our eCLIP experiments was manifest with the considerable overlap of significantly enriched peaks from two biological replicates (Extended Data Fig. 10c). After filtering those clusters according to their binding affinity to realize quality control, we noticed DDX43 displayed higher enrichments on the 3′ UTR (Extended Data Fig. 10d), and most preferable binding targets of DDX43 located at UTR regions (5′ UTR and 3′ UTR) (Extended Data Fig. 10e). These data support that RNA velocity analysis using 3′ end 10x Genomics scRNA-seq data could reflect the cell differentiation deficiency caused by 3′ UTR-bounding DDX43.

To determine the key perturbated phase in STids, we need to conduct differential expression gene (DEG) analysis between perturbated samples and wild-type samples across four sub-clusters. Using correlation of Fold Changes as the metric to evaluate the DEG consistence of two replicates (Method), we found the two replicates of *Ddx43^KI/KI^* show better consistence compare with the two replicates of *Ddx43^−/−^* (Extended Data Fig. 11a, b), suggesting that the DEG results between *Ddx43^KI/KI^* and wild-type samples are more reliable. Therefore, we zoomed the perturbation effects of knock-in samples. We overlapped DDX43 binding targets with differential expression genes (FDR-corrected p value < 0.05) for each sub-cluster, and denoted those genes as differential expression targets (DETs). Interestingly, we found the (DETs) in sub-cluster 0 showed much higher expression difference (abs(avg(knock-in sample) – avg(wildtype))) than other sub-clusters (Fig. 5i). We also found that the expression difference of all binding targets of DDX43 in sub-cluster 0 still showed high expression difference (Extended Data Fig. 12a). Meanwhile, sub-cluster 1 owned the vastest number of DETs (Fig. 5i, k). Furthermore, we discovered that most of DEGs in the sub-cluster 0 and sub-cluster 1 were down-regulated (Fig. 5j, 82.8% in sub-cluster 0 and 68.7% in sub-cluster 1), consistently, most of DEGs in sub-cluster 1 of *Ddx43^−/−^* sample were also downregulated (Extended Data Fig. 12b). In addition, the binding targets of DDX43 showed significant enrichments (P-value < 0.001, Fisher exact test) in those downregulated genes of sub-cluster 0 and 1 (Fig. 5k). These results pointed to a central role of DDX43 at the early stage of STids.

### Dynamical network analysis identifies *Elfn2* as a hub gene regulated by DDX43

We proceeded to elucidate the molecular dynamic network related to the DDX43 regulation in the early stage of STids. We analyzed the gene expression covariation of the down regulated DEGs in sub-cluster 0 and 1 from *Ddx43^KI/KI^* samples through WGCNA (weighted gene co-expression network analysis)^73^, and observed four major categories of transcriptional gene modules in characterized patterns (Fig. 6a). We also quantified those modules activity by AUCell and fitted a continuous curve alongside the velocity pseudo-time with local polynomial regression. We noticed most of the modules were gradually upregulated in activity and were largely involved the biological processes such as nucleus organization, fertilization, and mitochondrial respiratory chain complex assembly, corresponding to the functions in the later stage of STids, indicating DDX43 regulates genes enriched and highly expressed in later stage modules at the early stage. Strikingly, we noted there existed a co-expression module (donated as brown module) strongly altered at early stage, with the activity was consistently inhibited in *Ddx43^KI/KI^* cells. The genes in this module were largely involved with chromosome organization and DNA conformation process. Consistantly, the analyses results of weighted gene co-expression network using the down-regulated genes of sub-cluster1 in *Ddx43^−/−^* samples also showed a co-expression module (denoted as green module) strongly altered at early stage, with the activity was also consistently inhibited in *Ddx43^−/−^* samples (Extended Data Fig. 12c, d). This module also enriched with the pathways like DNA conformation, histone modification and chromatin remodeling (Extended Data Fig. 12e). Suggesting that the dysfunction of epigenetic regulation in early stage of STids also act important role in *Ddx43^−/−^* samples. Combining the dysfunctions in H4K8ac eviction (Fig. 3f), DNA repair (Fig. 3g and Extended Data Fig. 5b), and TNP-to-PRM transition (Fig. 3a, b, c, d, e) in previous results and activation of the epigenetic modification associated module in the early-stage STids (Fig. 6a and Extended Data Fig. 12c, d, e), we hypothesized regulation of this module may act as the origin events during spermatids differentiation

**Figure 6.**
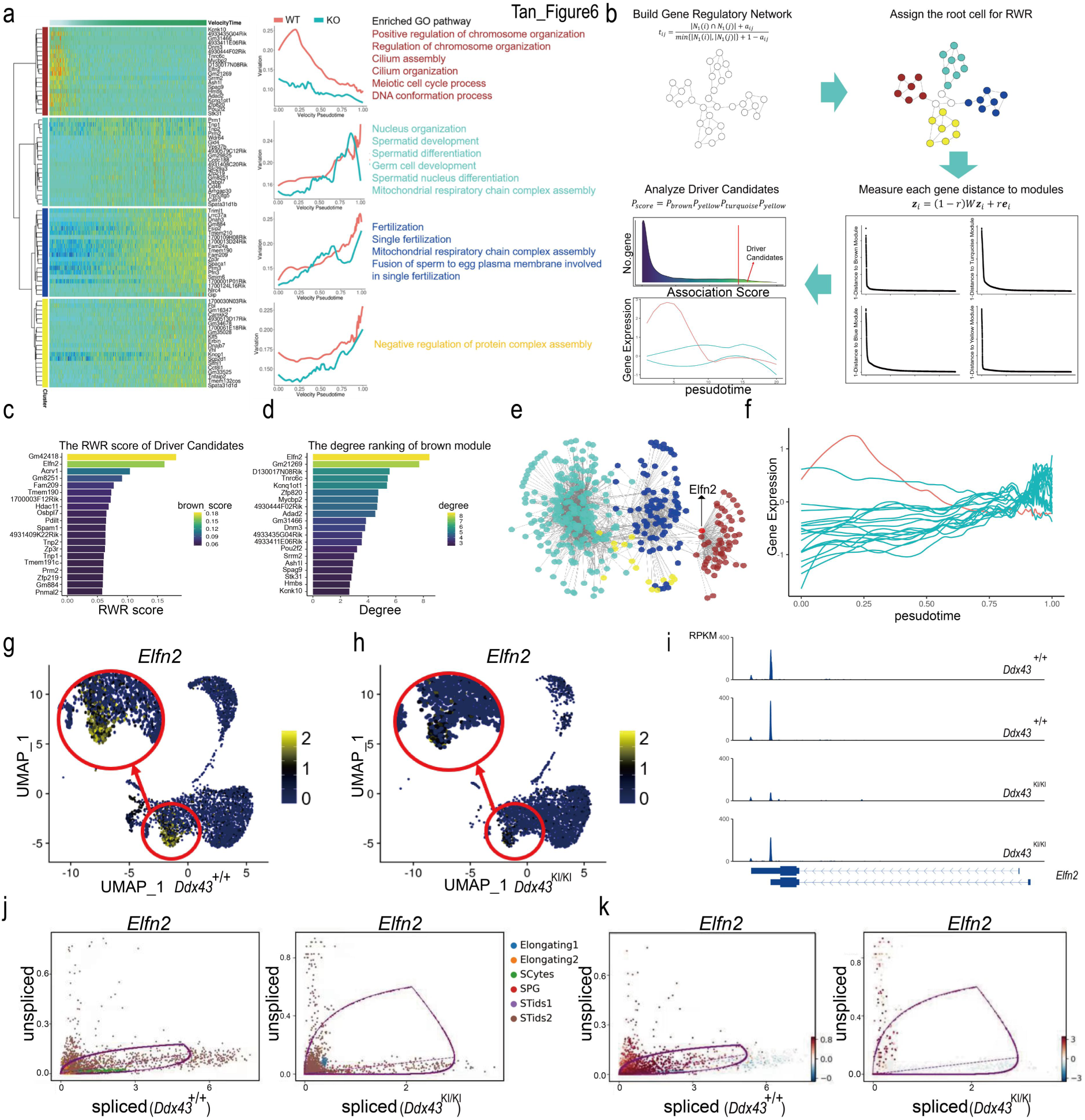
Dynamic network analyses for identifying DDX43-regulated driver genes. (a) WGCNA clustering of genes exhibiting down-regulated expression during subtype-0 and subtype-1. Each row represents a gene, and each column represents a single cell, with columns/cells placed in velocity pseudotime order and depicted by a thick colored line (top). Gene expression levels utilize a Z score transformation. (b) A workflow of page ranking algorithm to identify regulation driver candidates. In step 1, we converted the topological overlap matrix to markov transition matrix through row normalization. In step 2, we assigned the diffusion root through regarding those cells belong to gene co-expression module as the seed cells. In step 3, we performed message passing and got four rank score, each measure the proximity of each gene to the module. In final step, we aggregated those scores and identified the overall highly ranked genes as the driver candidates. (c) Barplot of association rank of driver candidates to the brown module, each bar is colored according to each gene topological associate score to brown module. The score is calculated from random walk with restart algorithm in panel b. (d) Barplot of the network connectivity of driver candidates. Each bar is colored according to each gene connectivity in the network, which reflects the gene’s importance to the biological systems (e) The gene regulatory network related to the dysregulation in subtype-0 and subtype-1. Each node represents a gene colored according to the module it belongs to. (f) Expression levels of driver candidate genes during spermatids developments. The x-axis represents velocity pseudotime, and the y-axis represents gene expression levels. The expression curve of *Elfn2* is marked in red. (g, h) Expression patterns of *Elfn2* in *Ddx43^+/+^* (g) and *Ddx43*^KI/KI^ (h) spermatogenic cells, with their expression projected onto the UMAP plot. (i) The track plot of *Elfn2* in *Ddx43^+/+^* and *Ddx43*^KI/KI^ subtype-0 cells, in two paired replicates. (j, k) The gene life cycle of *Elfn2* in *Ddx43^+/+^* (j) and *Ddx43*^KI/KI^ mutant (k) cells. The x-axis represents the spliced RNA expression level, the y-axis represents the unspliced RNA expression level. Each dot represents a cell colored according to the cell types (left panel) or velocity values (right panel).

We thus attempted to verify this assumption. To investigate the core genes related to the dysfunction of this brown module, we used page ranking algorithms on weighted gene-gene co-expression network, coupled with their topological distance to the four modules, to identify the top genes that proximal to the four modules (Fig. 6b). We noticed some genes of the top-gene list are associated with spermiogenesis (Fig. 6c, d), like *Acrv1* (an evolutionary conserved acrosomal matrix protein), and three core genes engage in TNP-to-PRM transition (*Tnp1*, *Tnp2*, *Prm1*). Also, we payed attention to some testis-unique or -high expression genes regardless of their so far limited functional clues for spermatogenesis, such as *Elfn2*, *Fam209*, *1700003F12Rik*, and so on. Considering the relatively late influence on genes required for TNP-to-PRM transition, *Tnp1*, *Tnp2*, or *Prm1* is less likely to be the origin of the *Ddx43^KI/KI^* defects. Therefore, we pointed to *Elfn2*, known as an epigenetic regulator highly expressed in the nervous system and testis^74, 75^. *Elfn2* is not only ranked as the top 2nd gene close to the ‘brown module’ (Fig. 6c), but also identified as one hub gene which owns the highest connectivity compared with other partner genes in ‘brown module’ (Fig. 6d, e). *Elfn2* is also identified as the top hub gene in epigenetic modification related module (green module) in *Ddx43^−/−^* sample (Extended Data Fig. 12f). What’s more, *Elfn2,* expressed at the early stage of spermatids (Fig. 6d, f),is the top down regulated DETs of DDX43. The gene expression analysis also assisted to determine the dysregulation of *Elfn2* in the early stage of STids in *Ddx43^KI/KI^* and *Ddx43^−/−^* testes (Fig. 6g, h, Extended Data Fig. 12g). In addition, the track plot further confirmed the exist of the *Elfn2* transcript in STids sub-cluster 0 (Fig. 6i). Gene spliced-unspliced life cycle analysis also indicated that *Elfn2* tends to have higher ratio of unspliced transcripts in *Ddx43^KI/KI^* testis (Fig. 6j, k). Taken together, our integrated analyses suggest that the downregulated *Elfn2* may lead the problematic translation of *Elfn2* mature protein and further the imbalance of the ‘brown module’ network.

### Knockdown of endogenous *Elfn2* recapitulates deficiencies from *Ddx43* mutant

To directly assess whether endogenous *Elfn2* contributes to chromatin remodeling during spermiogenesis, we first examined its testicular protein levels at four postnatal time points. Early at P21, concomitant with the appearance of round spermatids, ELFN2 protein abundance was equally low in wild-type versus *Ddx43^KI/KI^* and *Ddx43*^−/−^ mutant testes (Fig. 7a). At P28, P35 and P60, when round spermatids accumulated, ELFN2 protein levels increased but were significantly decreased in mutant testes, compared to wild type (Fig. 7a). These results are in concert with our scRNA-seq analytical data showing downregulation of *Elfn2* mRNA, as well as our observation of the phenotypic defects emerging evident from P28 (Extended Data Fig. 2e). To elucidate the *in vivo* function of *Elfn2*, we knocked down *Elfn2* through testis microinjection of antisense oligonucleotides (ASOs), as we established recently^76^. We validated knockdown efficiency 11 days after ASO transduction to P35 testes by western blot analysis. ELFN2 protein levels declined in ASO knockdown versus control (Fig. 7b), proving validity of our *Elfn2* knockdown model. Histological analysis of P35 testes revealed similar phenotypic defects in *Elfn2* knockdown seminiferous tubules compared with *Ddx43* mutant mice (Fig. 7c). Meanwhile, we co-immunolabeled γH2AX and PNA, and observed strong γH2AX localization on step 14 spermatids in both *Ddx43* mutant and *Elfn2* knockdown seminiferous tubules at stage II, but not in *Ddx43^+/+^* and *Elfn2* controls (Fig. 7d). In parallel, co-immunolabeling H4K8ac and PNA showed presence of H4K8ac-positive step 14 spermatids only in *Ddx43* mutant and *Elfn2* knockdown testes (Fig. 7e). These images displaying abnormally remained γH2AX and H4K8ac in P35 testes largely resemble those described in P60 testes (Fig. 3f, g). To further address how DDX43 regulates *Elfn2* expression, we employed dual luciferase reporter assay in HEK293 cells, where we introduced a sequence of *Elfn2* 5′ UTR into the reporter vector and overexpressed exogenous DDX43 protein. As a result, DDX43 overexpression weakened the reporter gene translation (Extended Data Fig. 13a), suggesting that DDX43 may act upon *Elfn2* 5′ UTR. In addition, using this overexpression model, we validated that DDX43 binds *Elfn2* transcript by RNA immunoprecipitation (RIP) followed by qRT-PCR (Extended Data Fig. 13b). These results elucidate that DDX43-mediated regulation of *Elfn2* expression is at least partly responsible for maintaining the normal process of chromatin remodeling during spermiogenesis.

**Figure 7.**
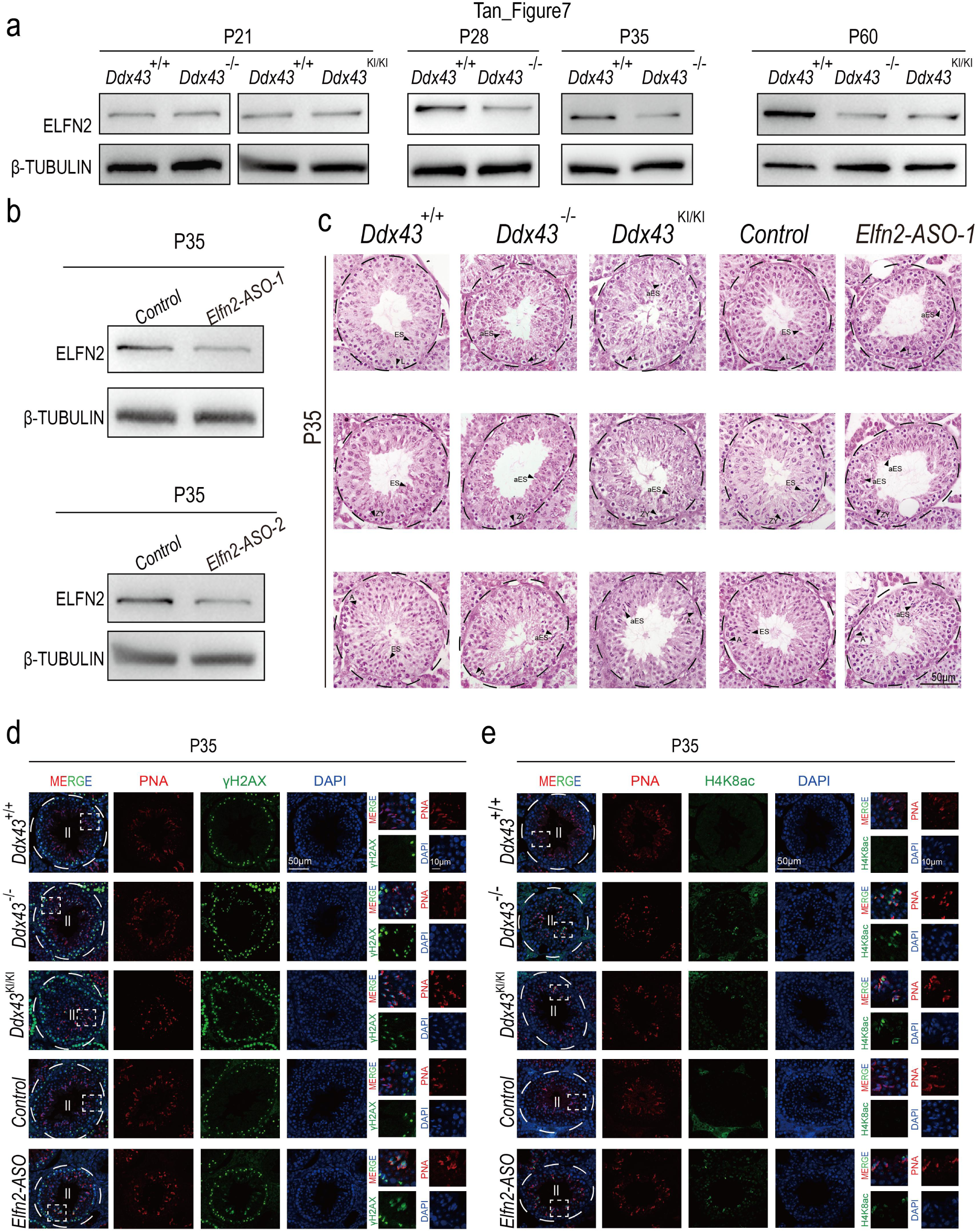
Endogenous knockdown of DDX43-targeted gene *Elfn2*. (a) Western blot analyses of ELFN2 protein in *Ddx43^+/+^* and *Ddx43* mutant testes at indicated postnatal time points. (b) Western blot analyses confirmed ELFN2 protein was efficient knockdown in *Elfn2* knockdown mice lysates. β-TUBULIN serves as an internal loading control. (c) H&E staining analyses using P35 testes sections prepared from *Ddx43^+/+^*, *Ddx43*^KI/KI^, *Ddx43^−/−^*, *Elfn2* control and ASO-transduced P35 testes. Scale bar is indicated. (d, e) Immunofluorescence analyses using P35 testes sections of indicated genotypes with two combinations: γH2AX (green) and PNA (red) (d); H4K8ac (green) and PNA (red) (e). DNA was counterstained with DAPI. Stage numbers and scale bars are indicated.

## Discussion

It’s well-documented that post-meiotic germ cells undergo chromatin remodeling involving multiple intricated events that are chronologically coordinated, accompanied by progressive compaction of sperm nucleus. scRNA-seq have provided delicate new insights into how spermatids development goes through a highly dynamic process of transcriptome transformations. Thereupon, it’s tempting to ask whether there exists a factor that bridges regulation of developmentally dynamic transcriptome and shaping of chromatin architecture during spermiogenesis. Such a regulator could be an RNA-binding protein that should act upon meiotic exit, but so far has yet to be identified. In the present study, we show that the RNA helicase DDX43 is expressed in a spatiotemporal fashion and its protein levels prominently span late meiotic and early post-meiotic stages. Genetic mutation of *Ddx43* renders minimum disturbance to meiosis, whereas leads to severe multistep defects in developing spermatids, failure to produce mature spermatozoa, and ultimately male infertility. Importantly, *Ddx43*-deficient spermatids display aberrant occurrence of stage-dependent events of chromatin remodeling including H4K8ac and γH2AX eviction and TNP-to-PRM substitution. The reasons that these phenotypic and molecular outcomes stem from an unprecise control of chromatin remodeling events are two-fold. First, despite that partial mutant spermatids may be developmentally retarded or not purged from prior stages, the overall heterogeneity in molecular defects present at same steps precludes a fully uniform arrest unreachable to its gene expression steps and thus points to protein behavioral abnormality. Second, our scRNA-seq unravels the dysregulation of expression of a group of enriched and related genes in mutant testes, and therein identified *Elfn2* as a targeted hub gene downstream of DDX43 at early-step spermatids. Therefore, our work demonstrates an important role of DDX43 and highlights its stage-specific transcriptome control in orchestrating chromatin remodeling.

To seek the molecular function of DDX43 *in vivo*, we explored three possibilities before employing scRNA-seq. First, we created gene-edited mice bearing a point mutation in the ATP hydrolysis site of DDX43. This site is highly conserved and validated for functional importance^50, 51, 77^. Regrettably, DDX43 protein itself is almost absent in this mutant testis. Similar outcomes of point mutation-elicited self-depletion are attributed to protein destabilization^78, 79^. Although it is difficult to distinguish the bona fide effect between abolishing helicase activity and loss of the majority of DDX43 protein per se, it is still worth analyzing for mechanism decipher of DDX43 protein. Second, we examined the effect of loss of DDX43 on the piRNA pathway wherein no difference is found in piRNA abundance, Piwi localization or retrotransposon derepression. We previously studied two distinct knockout mouse models of another testis-specific RNA helicase MOV10L1: one displays de-repression of retrotransposon and meiotic arrest^43^; the other exhibits blockade of pachytene piRNA biogenesis and spermiogenic arrest^64^. As such, two different RNA helicases herein with overlapping expression are not committed to a joint pathway. Third, to study DDX43 as an RNA regulator, we performed RNA-seq analysis using postnatal testes enriched for certain types of germ cell. Unexpectedly, *Ddx43* mutant does not cause a major change in gene expression. Some other studies for chromatin remodelers in spermatids also show limited power of the bulk RNA-seq in revealing an overall alteration in gene expression^6, 16^. The challenges posed by all negative results represent opportunities to envision an unusual mode of action of DDX43 in testis. As a scenario for our case, within a certain narrow developmental window may DDX43 control the abundance of determinant genes, whose expression fluctuations, nevertheless, are unable to be manifested into significant values of RNA-seq. We thus proceeded to subject *Ddx43* mutant model to scRNA-seq to deeply analyze transcriptome pattern-specified populations of wild-type verse mutant germ cells.

We recently developed a competitive tool, termed ICAnet, which leverage independent component analysis and biological network (e.g. protein-protein interaction network) to extract meaningful signal from single cell transcriptome data. Therefore, ICAnet could give a better single cell clusters, and even identify rare cell types from the dataset^35^. We herein fed this method into our *Ddx43* mutant mouse model and generated an atlas of DDX43-regulated transcriptomes with full coverage of major male germ cell populations during mouse spermatogenesis. The substantial proportion of differently grouped spermatid cells, which arise from a small proportion of spermatogonia and spermatocyte cells, provide a typical test case in which the direction of spermiogenic differentiation can be validated by lineage tracing. RNA velocity-based approach of trajectory inference on using single-cell transcriptomic data has recently been adopted to visualize the dynamic movement of individual cells in various biological processes, with no demand of prior knowledge^30, 80, 81^. Recently, a new computational tool named scVelo, using Expectation and Maximization (EM) algorithm to infer full dynamics of splicing kinetic, generalizes RNA velocity estimation to system with transient cell states. To infer intrinsic developmental trajectory of spermatogenesis on the basis of our aforementioned dissected germline subpopulations, we adapted scVelo into our scRNA-seq analytic pipeline to study our mouse models. Using velocity-directed Markov chain simulation to predict the differentiation terminal probability along the spermatogenesis, we observed that RNA velocity of wide-type germ cells showing a strong directional flow towards the termini in Elongating Cell (ES). However, mutant germ cells exhibit a new differentiation terminal point in spermatids, which prompted us to an in-depth analysis of the transcriptomic perturbation in spermatids. Our study has exemplified a paradigm where RNA velocity can be used to depict gene-regulated dynamic processes in spermatogenesis, and will greatly facilitate trajectory inference of germ cells under various genetical contexts.

Although scRNA-seq could help us identify key sub-clusters of cells and the genes that change in those cells, it was still not enough to determine the initially affected cell populations and the changed genes that are more likely to be “causative”, i.e., directly targeted and mediate subsequent chain reactions. eCLIP-seq provides us a series of candidate targets that are, in principle, bound by RNA-binding protein. We thus performed eCLIP-seq to acquire RNA substrates of DDX43. Combinatorial analysis of scRNA-seq and eCLIP-seq data enables us to narrow down the list of genes that are defined to be DDX43-bound and differentially expressed genes derived from the core cell clusters in spermatids. Remarkably, the binding targets of DDX43 were significantly enriched in those downregulated genes in the sub-clusters 0 and 1, confirming our prediction that the early post-meiotic cells may be the initial populations controlled by DDX43. Coupling eCLIP-seq that is specialized for RNA-protein interactions in testis with scRNA-seq that unveils regulation of target modules will be an impetus towards ultimately bringing them together at single-cell scales, enlightened by recent advances in CLIP technologies^82–84^.

Weighted correlation network analysis (WGCNA) can be used to screen gene network for identifying candidate hub gene from clusters constituted by highly correlated genes, and numerous biological contexts have been successfully addressed by these methods^85^. This algorithm enables us to measure the gene topological properties and the summarized gene co-expression module activity helps us to relate modules and genes to external sample/cell traits (genotype/phenotype)^73, 86, 87^. In this study, we fed WGCNA into our analysis to build a gene co-expression network for the downregulated genes in the early stage (sub-cluster 0 and 1) of spermatids and quarried four gene modules. Using AUCell and local polynomial regression, we fitted a continuous curve to describe the dynamical changes of the expression activity of those four modules alongside the velocity pseudo-time. We found that the brown module strongly altered at the early stages, and the genes function in this module happened to be echoed with the observed dysfunctions in chromatin remodeling of *Ddx43* mutants. We thus further applied page ranking algorithms on weighted gene-gene co-expression network and tapped out that *Elfn2* may be the key target of DDX43 and function in spermiogenesis. The following validation experiments confirmed our inference. In detail, the *Elfn2* knockdown spermatids exhibit mutant-comparable defects, regardless that *in vivo* knockdown does not perfectly phenocopy knockout. Differing from the consequence of *Elfn2* knockdown, loss of its upstream regulator DDX43 yields a pleiotropic outcome from impacting a catalog of molecular pathways. These analyses have enabled effective identification of novel gene expression modules, and more importantly, key target genes that are regulated by an RNA-binding protein. Future studies by probing germline stage-specific interaction of *Elfn2* mRNA with DDX43 protein as well as creating *Elfn2* conditional knockout mouse model would be an extension to elucidate when and how *Elfn2* mediates the DDX43-orchestrated process of chromatin remodeling.

Currently, scRNA-seq is being utilized to delineate cellular and molecular processes with response to various biological or environmental perturbations^88–90^. As a unique study presented herein is our scRNA-seq-based investigation of single gene-governed spermatogenesis. Our study defines an unexpected RNA regulatory pathway of DDX43 in leading the progressive differentiation trajectory of germ cells. As a framework, the single-cell revolution of scRNA-seq, integrated with sophisticated analytical methodologies, genetic models and molecular approaches, will dramatically elevate our ability to resolve the genetic regulation of the highly ordered dynamics of spermatogenesis.

## Materials and Methods

### Animals

Gene locus *Ddx43* is located on mouse chromosome 9 and comprises 17exons (Fig. 1g). For generating knockout mouse model (termed *Ddx43^−/−^*), we designed two guide RNAs (gRNA) targeted exon 4. Briefly, sgRNAs sequences were cloned into pUC57-sgRNA vector (Addgene 51132). sgRNAs were transcribed using the MEGA shortscript T7 Transcription Kit (Life technologies, AM1354) and purified with RNeasy Mini Kit (Qiagen, 74104). sgRNAs (5 ng/μl each) and Cas9 mRNA (20 ng/μl, Thermo Fischer Scientific, A29378) were introduced into C57BL/6J mouse zygotes by electroporation. Founder mice were selected according to Sanger sequencing results and crossed with wild-type C57Bl6/J (Jackson) partners to obtain germline transmission. All animals were handled as stipulated by the Guide for the Care and Use of Laboratory Animals at Nanjing Medical University, and raised in a suitable environment with moderate light, clean water and food. Regardless of the experiment, all mice were selected at random without considering any subjective factors, and at least 3 animal samples in each experiment. Using analogous methods, we also created knockin mice with a point mutation in the DDX43 ATPase motif (DEAD-DQAD).

### Antibodies

#### Commercial Antibodies

The following antibodies were purchased: anti-SYCP3 (Abcam, ab97672, IF:1:100), anti-SYCP3 (Abcam, ab15093, IF:1:100), anti-SYCP1 (Abcam, ab15090, IF:1:100), anti-γH2AX (MerckMillipore, 16-202A, IF:1:800), anti-TNP2 (Santa Cruz, sc-393843, IF:1:50), anti-PRM2 (Briar Patch Biosciences, Hup 2B, IF:1:300), anti-H4K8ac (ABclonal, A7258, IF:1:500), anti-ELFN2 (Novus Biologicals, 90569, WB: 1:1000), anti-H3 (Abcam, ab1791, WB: 1:2000), anti-GAPDH (Abcam, ab8245, WB: 1:10000), anti-TUBLIN (Sigma, T8328, WB: 1:2000). For immunofluorescence studies the following secondary antibodies were used: anti-rabbit (Jackson ImmunoResearch 488, 711225152, IF: 1:350), anti-rabbit (Jackson ImmunoResearch 594, 711585152, IF: 1:350), anti-mouse (Jackson ImmunoResearch 488, 715545150, IF: 1:350). For western blot analyses the following secondary antibodies conjugated to Horse Radish Peroxidase were used: anti-rabbit IgG HRP-linked (ABclonal, AS014, WB: 1:5000), anti-mouse IgG HRP-linked (ABclonal, AS003, WB: 1:5000).

#### Other antibodies

Goat-ACRV1 is a kind gift from Eugene Yujun Xu lab in Nanjing Medical University. Custom-made DDX43 antibody is provided by ABclonal. Briefly, a cDNA fragment of *Ddx43* corresponding to amino acids 630-646 (EMEKKMGRPQGKPQKFY) was cloned into the pGEX-4T-1 vector. Using glutathione-sepharose beads, the GST-DDX43 (630-646) fusion protein was purified from Rosetta bacteria. Purified recombinant DDX43 protein was next administered to two rabbits. DDX43 antiserum was collected and affinity-purified. Finally, we proved effectiveness of the purified DDX43 antibody in western blot (1:1000), immunostaining (1:500), and CLIP experiments (10 µg).

### Western blot

We first collected tissues, quickly washed twice in cold PBS, and then homogenized with a glass tissue homogenizer in lysis buffer (50 mM Tris pH 7.5, 150mM NaCl, 1% NP-40, 0.5% sodium deoxycholate) containing protease inhibitors (Roche, 11697498001). Supernatant fractions were collected by centrifugation and boiled with 2× Laemmli sample buffer (Bio-Rad, 1610737) at 95 °C for 8-10 min. The proteins were separated by Sodium Dodecyl Sulfate-Polyacrylamide Gel Electrophoresis (SDS-PAGE, 10% acrylamide running gel), and then electrically transferred from Gel to 0.45 µm polyvinylidene fluoride (PVDF) membranes (Bio-Rad, 1620184) at 90 V for 30 min, plus 120 V for 90 min. After washing membranes with Tris-buffered saline (TBS, 20 mM Tris, 150 mM NaCl, pH 7.6), membranes were blocked with 5% skimmed milk in TBST buffer (TBS with 0.05% Tween20) at room temperature for 1 h and then incubated with antibodies overnight. The next day, membranes were washed with TBST buffer for 5 min and incubated with HRP conjugated secondary antibodies at room temperature for 1 h. Signals were visualized by an enhanced chemiluminescence detection system (Tanon-5200).

### Quantitative and semi-quantitative RT-PCR

Total RNA was extracted from tissues or cells with Trizol reagent (Thermo Fisher Scientific, 15596026) and DNaseI (Amp grade 1.5 µ/µl, Invitrogen RNase free). Prime Script RT Master Mix (Takara, RR036A) was used to reverse transcribe 1 µg of total RNA into cDNA. SYBR Green Premix Ex Taq II (Takara, RR820A) was used to analyze the quantitative RT-PCR (RT-qPCR) with 2 µl diluted cDNA as template. *Rplp0* (also known as *36b4*) served as internal control. For semi-quantitative PCR, 1µl diluted cDNA was used as the template for each reaction with Emerald Amp GT PCR Master Mix (Takara, RR310). After appropriate amplification cycles, half of the reaction products were analyzed by 1% agarose gel electrophoresis. *Arbp* was used as loading control, and all primer sequences were listed in Table. S1.

### Histology and transmission electron microscopy

To prepare paraffin sections, testes and epididymis were fixed in 5 ml Bouin’s solution (Sigma, HT10132) overnight. The next day, tissues were dehydrated with a graded series of ethanol (70%, 2h; 80%, 2h; 95%, 2h; 100%, 2h), soaked the tissues in xylene (Biosystems) for 3 h, replaced by paraffin for another incubation at 65 °C overnight, and then embedded into plastic molds with paraffin. Samples were sliced into 5 µm by microtome (Leica, RM2135). The sections were dried at 60 °C and then stored at room temperature. For histological analyses, the sections were deparaffinized, rehydrated, and then stained with Hematoxylin and Eosin (H&E, Sigma-Aldrich) or Periodic acid-Schiff (PAS) reagent. The sections were covered with coverslips immediately after a few drops of Neo-Mount (Merck) were deposited on them. For transmission electron microscopy analysis, we used 5% glutaraldehyde in 0.2 M cacodylate buffer to fix the tissues at 4 °C overnight. The next day, tissues were washed with 0.2 M cacodylate buffer, dehydrated in increasing concentration of ethanol, embedded and polymerized by the automated microwave tissue processor (Leica EMAMW). After that, samples were cut into small slices with the LEICA Ultracut UCT ultramicrotome (Leica Microsystems). Ultrathin sections were stained with uranyl acetate and lead citrate and examined by TEM (JEOL, JEM-1010).

### Immunofluorescence and chromosome spread

To prepare frozen sections for immunofluorescence analysis, testes were fixed in 10 ml 4% paraformaldehyde (PFA) at 4 °C overnight. The next day, tissues were dehydrated with 15% sucrose in 1x PBS until totally sunk to the bottom of the tube, transferred to 30% sucrose for further dehydration at 4 °C overnight. And then, embedded in OCT (Fisher Scientific, 14-373-65) and cut into 5 µm in cryostat (Thermo Scientific Cryotome FSE). Slices were stored at -80 °C. For immunofluorescence analysis, the slices were treated as follows: air dried at room temperature, washed twice with PBS, permeabilized with 0.5% Triton X-100 in PBS at room temperature for 10 min. After washing three times with PBS, samples were blocked with 5% serum in TBS-T (TBS, 20 mM Tris, 150 mM NaCl, pH 7.6, and 0.1% Tween20) at room temperature for 1 h, incubated with primary antibodies in suitable concentration at 4 °C overnight, and then conjugated with secondary antibody and DAPI at 37 °C for 1 h. Chromosome spreads of prophase I spermatocytes were conducted as we performed previously^27^. All fluorescence images were taken by Carl Zeiss LSM800 confocal microscope.

### Isolation of spermatogenic cells

Mouse spermatogonia, pachytene spermatocytes, round spermatids and elongated spermatids were isolated using a BSA gradient method as previously described^21^, with minor modifications. In brief, after euthanasia, mice testes were harvested, quickly washed twice in cold PBS, and then digested in 10 ml DMEM (Gibco, 12800017) containing collagenase (1 mg/ml, Gibco, 17100-017) for at 37 °C 15 min with gentle shaking. The dispersed seminiferous tubules were collected by spinning at 300 ×g at 4 °C for 5 min, and washed twice with DMEM. Tubules were then digested in 45 ml 0.25% Trypsin (Gibco, 25200-072) containing DNase I (1 mg/ml, Qiagen) at 37 °C for 5 min with gentle shaking, filtered through 40 µm Nylon Cell Strainer (BD Falcon, 352340) and resuspended with 25 ml DMEM containing 0.5% BSA. After loading the single-cell suspension into the separation apparatus (ProScience, Canada), germ cell populations were separated by sedimentation at unit gravity for 3 h through a gradient of 2-4% bovine serum albumin (BSA) solution. After that, cell fractions were harvested, and identified by their morphological characteristics. The purity of isolated SG, Spg, PS, and RS was approximately 85%, 90%, 90%, and 85%, respectively.

### Isolation of nuclear and cytoplasmic fractions

Subcellular fractions were extracted as we described previously^91^, with minor modifications. Briefly, about 100 mg testes tissues were homogenized in 1 ml Cytoplasmic Extraction Buffer (250 mM sucrose, 10 mM Tris-HCl, pH 8.0, 10 mM MgCl2, 1 mM EGTA, 1× protease inhibitor cocktail III) by pestling about 100 times. Nuclei were pelleted by spun down at 300 ×g at 4 °C for 5 min, and the supernatant was collected as the cytoplasmic fraction. The cytoplasmic fraction needs another high-speed centrifugation at 14000 ×g at 4 °C for 10 min to acquire purified cytoplasmic protein. The nuclear pellet was washed three times in Cytoplasmic Extraction Buffer plus once with PBS, dissolved in Nuclear Extraction Buffer (250 mM sucrose, 10 mM Tris-HCl pH 8.0, 10 mM MgCl2, 1 mM EGTA, 0.1% Triton X-100, 0.25% NP-40, and 1× protease inhibitor cocktail III), and then centrifuged at 14000 ×g for 15 min to collect supernatant as the nuclear protein.

### ASO-based knockdown in testis

We performed the transduction as previously described^76^. In brief, P24 mice were anesthetized by tri-bromoethanol. Testes on two sides were exteriorized in order through incisions on abdomen. Under a stereoscopic microscope, one testis was injected with 4 µl *Elfn2*-targeting ASO dilutions and the other was injected with 4 ul none-targeting ASO dilutions as control. After recovery, mice were fed as before. Eleven days later, all testes were harvested for the subsequent experiments. ASO oligonucleotide sequences were listed in Table. S1.

### Plasmid construction and Dual Luciferase Reporter assay

The *Elfn2* (ENSMUST00000088592.6) 5’UTR region and the full-length cDNA of *Ddx43* (*Ddx43^+/+^* and *Ddx43^−/−^*) were PCR amplified from mouse genomic DNA, then cloned them into psiCHECK™-2 luciferase vector and pRK5-FLAG vector, respectively. These restructured vectors are termed as *Elfn2*-5’UTR, *Ddx43* (+) and *Ddx43* (-). The primers for cloning were listed in Table S1. We co-expressed these restructured vectors in HEK293T cells. After transfection for 48 h, the cells were lysed and measured with Luciferase Assay Kit (Promega, E2920). Firefly luciferase values were normalized to Renilla and measurements of mean ± S.D of relative luciferase unit per microgram of protein were taken in triplicates and represented graphically as mean ±S.D on a MS-Excel sheet.

### Radiolabeling detection of piRNA and CLIP-captured RNA

Mouse testicular piRNA was radiolabeled as described previously^64^. Briefly, total RNA (1 µg) extracted from adult mouse testes was treated with alkaline phosphatase (NEB), de-phosphorylated, and 5’ end-labeled using T4 polynucleotide kinase (NEB) and [γ-^32^P] ATP. Then, we ran Urea-PAGE gel to separate the ^32^P labeled RNA. Finally, by exposing the gel to a phosphorimager screen and scanning with a Typhoon scanner (GE Healthcare), radioactive signals were visualized.

We performed DDX43 conventional CLIP essentially as previously for MOV10L1^91^. And the exhaustive details were described by Vourekas^92^. In brief, 100 mg testis tissues were detunicated with 4 ml cold PBS, UV-irradiated on a 10 cm plate at 254 nm three times inside UV cross-linker (UVP CL-10000), centrifuged at 1000 ×g, 4 °C for 1 min to collect the pellet, flash-frozen in liquid nitrogen, and then stored pellet at −80 °C or immediately used. To continue CLIP, pellets were dissolved with 300 µl 1X PMPG buffer (1x PBS no Mg2+/Ca2+, 2% Empigen) containing protease inhibitors (Roche, 11873580001) and RNasin (Promega), treated with DNase (Promega, M6101) at 37 °C for 10 min in a Thermomixer, and then spun down at 15,000 rpm at 4 °C for 30 min. The supernatant was then immunoprecipitated with anti-DDX43 antibodies using protein A Dynabeads (Invitrogen, 10002D). Beads were washed with 1x and 5x PMPG and treated with alkaline phosphatase (NEB) to de-phosphorylate DDX43-crosslinked RNA on beads. And then, the [γ-^32^P] ATP-labeled RNA linker (RL3) was added to beads for RNA ligation overnight. The next day, after stringent wash steps, crosslinked DDX43 RNPs were boiled with NuPAGE™ LDS (Life Technology, NP0007), separated by NuPAGE™ 10% Bis-Tris (Life Technology, NP0323BOX), and transferred onto nitrocellulose (Invitrogen, LC2001). Membranes were exposed to phosphorimager screen for autoradiography.

### Enhanced CLIP (eCLIP)-seq

We prepared eCLIP libraries as previously described^71, 72^. Briefly, testicular seminiferous tubules were harvested and UV-irradiated as described for conventional CLIP, lysed and sonicated with 1ml fresh eCLIP lysis (50 mM Tris-HCl pH 7.4,100 mM NaCl, 1% NP-40 (Igepal, CA630), 0.1% SDS, 0.5% sodium deoxycholate (protect from light), 1:200 Protease Inhibitor Cocktail III), and then, incubated with RNase I (Life Technology, AM2295) and DNase (Promega, M6101) in a Thermomixer at 1200 rpm to fragment RNA at 37 °C for 10 min . Then, DDX43-RNA complexes were immunoprecipitated with anti-DDX43 antibodies, followed by dephosphorylation of RNA fragments and ligation of 3’ RNA adapter. After stringent wash with low- and high-salt buffers, eCLIP samples, as well as SMInput samples (as control library to subtract noise from eCLIP-seq library), were boiled from beads, run on an SDS-PAGE gel and transferred to nitrocellulose membranes. In parallel, a small proportion of samples was reserved for western blot analysis, which validates specific enrichment of DDX43 protein and identifies cutting territory on membrane. And then, RNA from both eCLIP and SMInput were harvested from the membrane by digesting the protein with proteinase K and Urea. The harvested RNA was reverse transcribed with Affinity Script (Agilent,600107), purified by Exo-Sap-IT (Affymetrix, 78201) to remove unincorporated primer, and further ligated with 3’DNA adaptor. Finally, we used Q5 PCR mix (NEB, M0492L) to amplify the libraries for high-throughput sequencing in Illumina platform.

### RNA-seq Libraries

We used TruSeq Stranded Total RNA kit (Illumina) for library preparation. Libraries were assessed with the Qubit and Tapestation for molarity and quality before submitting to the Illumina Hiseq X ten system.

### Single cell RNA-seq (scRNA-seq) Libraries

We performed testis scRNA-seq from adult *Ddx43^+/+^* and *Ddx43^−/−^* mice as we previously described^35^. In brief, testes were digested into single cell suspensions as the performance in isolation of spermatogenic cells. Cell concentration was counted by Cellometer Mimi (Nexcelom Bioscience). Mixed the cell suspension with Trypan Blue (Life Technology), and then counted the viability of cells by the introverted microscope. Only samples with >80% viability were loaded into Single Cell A Chip (10X Genomics, Chromium) and recovered about 5000 cells. Chromium Single Cell Controller (10X Genomics) was used to acquire single cell Gel Bead-In-EMusions (GEMs). Finally, libraries were constructed with Single Cell 3ʹ Reagent Kits v2 before sequencing by Illumina Novaseq.

### Bioinformatics analyses

#### Processing of RNA-seq Data

The reads were mapped to the GENCODE GRCm38 v23 transcript set using Bowtie2 and the gene expression level was estimated using RSEM.TMM (trimmed mean of M-values) was used to normalize the gene expression. Differentially expressed genes were identified using the edgeR program. Genes showing altered expression with p < 0.05 and more than 1.5 fold changes were considered differentially expressed.

#### Processing of eCLIP-seq Data

Reads were adaptor trimmed through cutadapt, and mapped to mouse genome (with repetitive elements-mapping reads discarded) with STAR. PCR duplicate reads were removed. We then used CLIPper to perform peak-calling. Input normalization of peaks was performed by counting reads mapping to CLIPper-identified peaks in eCLIP and paired SMInput dataset, with significance thresholds of *P* ≤ 10^-5^ and fold enrichment ≥ 4. The binding affinity of each peak is defined by counting reads mapping to peaks and perform TPM normalization.

### Processing of 10X Genomics single cell RNA-seq Data

#### Read filtering and alignment

Raw sequencing data were converted to fastq format using cellranger mkfastq (10x Genomics, v3.0.0). scRNA-seq reads were alighed to the GRCm38 (mm10) reference genome and quantified using cellranger count (10x Genomics, v.3.0.0). After collecting digital gene expression count matrix, we performed quality control to the cells based on the distribution of genes detected and UMIs, mitochondrial transcripts percent of each cell for all experiments. On the basis of these distribution, we filtered out the cells with detected genes ≤ 2.5 quantile or ≥ 97.5 quantile regarded as the outlier cells for each experiment, and we also filtered out cells with ≥ 10% of transcripts corresponding to mitochondria-encoded genes. Furthermore, we computed pearson residuals of our expression dataset through *SCTransforms* functions in Seurat, with regarding cell cycling score, mitochondria-encoded genes transcripts level, and the total number of UMIs for each cell as the covariates.

### Module Activity Matrix Construction and Cell Type Annotation

We used previously developed algorithms ICAnet^35^ to convert normalized gene expression matrix to the module activity matrix. ICAnet used both independent component and protein-protein interaction network to identify gene co-expression modules, using those modules could facilitate cell clustering and batch integrations. For WT samples, we used JADE algorithms to compute independent components combine with STRING^93^ network to identify modules through random walk trapping. Using identified module lists, we could quantify those modules activity on both WT and KO samples through AUCell^65^. We finally identified 1085 modules in our data, and used AUCell induced module activity matrix to perform downstream analyses. To define high quality reference atlas, we first analysis WT samples, and used their module activity matrix to infer top 20 principal components to calculate Euclidean distance among all pairs of cells and for each, identified its 30 nearest neighbors to define cell *k*-nearest neighborhood (KNN) graph. It then used the Louvain method of clustering using Jaccard distance among cells as weights, where the Jaccard distance between ant two cells was defined as the degree of overlap of their 30 nearest neighbors, then using UMAP^94^ to perform visualization. We then annotate each cluster according to known spermatogenesis marker genes expression pattern within each cluster.

### Mutant to Reference Mapping and Label transferring

After defining the reference map. We then using anchors base integration methods provided in Seurat. Using defined new assay, module activity matrix, we projected cells from mutant samples on reference samples through looking the mutual nearest neighbor cell in reference samples for each cell in mutant sample, and defined as anchor, we then transferred the labels (cell type annotations) in reference samples to the cells in mutant samples through those anchors, and only preserve those cells which prediction score ≥ 0.95 to get a high confidence prediction.

### Cell Differentiation Potency Quantification

To measure the cell differentiation potency, we used a signaling entropy model names single cell entropy (SCENT)^66^ to estimate each cell’s plasticity. SCENT could quantify the degree of uncertainty of a cell’s gene expression levels in the context of a cellular interaction network through constructing a stochastic matrix base on PPI network for each cell. However, the original signaling entropy score need to infer an invariant measure Π for each cell, which rely on singular value decomposition and it need to cost tremendous time for computation. Therefore, we used a global mean field approximation of the signaling entropy, that is, the pearson correlation of the cell’s transcriptome and the connectome from the PPI network.

### RNA velocity and Trajectory Analysis

To infer furture states of individual cells, we used spliced and unspliced transcript ratio to predict cell differentiation trajectory. This combinational analysis was performed using a combination of the Seurat and *scVelo*^36^ code. First, we used cellranger output aligned bam files as input for Velocyto to derive the counts of unspliced and spliced reads in loom format. Next, the non-spermatogenesis cells were excluded from the datasets, and we saved previously annotated cell type label, computed principal components and UMAP embedding (generated from “*integrate*” assay) and the spliced and unspliced read count matrix as new Seurat object, and converted the cleaned Seurat object into a .h5ad file for downstream analysis using python package scVelo. Each sample (*Ddx43^+/+^*, *Ddx43^KI/KI^* and *Ddx43^−/−^*) loom files were normalized and log transformed using scVelo functions normalize_per_cell() and log1p(), then we used to calculate first and second order moments for each cell across its nearest neighbors (scvelo.pp.moments(n_pcs = 20, n_neighbors = 30)). Next, the velocities were estimated and the velocity graph constructed using the scvelo.tl.velocity() with the mode set to ‘stochastic’ and scvelo.tl.velocity_graph() functions. Velocities were visualized on top of the previously calculated UMAP coordinates with the scvelo.tl.velocity_embedding() function. To compute the terminal state likelihood, the function scvelo.tl.terminal_states() with default parameters was used.

### Evaluating the confidence of DEG identification results

For each genotype, *Ddx43^KI/KI^* and *Ddx43^−/−^*, we have two biological replicates. Normally we performed DEG analysis through merging the cells from both samples, and using Wilcoxon rank-sum test to perform differential expression analysis for each gene. However, we need to account for the individual differences or batch effects may lead different DEG results. To assess the effects, we performed DEG analysis for each biological replicate to wild-type samples, and calculated the correlation of two gene expression FoldChange vector, which is derived from two biological replicate.

### Reclustering of STids and building PAGA graph

To identify the latent cell state within spermatids (STids). We extracted the cells from STids of integrated samples (includes all *Ddx43* genotypic samples), and re-computed the principal components (PCs). We used top 10 PCs to build cell KNN graph, and using Louvain clustering algorithm with resolution parameter set as 0.25 to identify cell sub-clusters. Using this pipeline, we found four sub-clusters within STids, denoted as sub-cluster 0-4. To assess the global connectivity topology between the Louvain clusters we appliced Partition-based graph abstraction (PAGA)^70^. We applied the tl.paga() function integrated in the Scanpy package to calculate connectivities and used the predefined Louvain clusters as partitions. The weighted edges represent a statistical measure of connectivity between the partitions. The orientation of the edges between each Sub-cluster is given by previous RNA velocity analysis.

#### Identification of DEGs

To identify the DEGs between *Ddx43^KI/KI^* and *Ddx43^+/+^* samples within each sub-cluster, we only extracted the normalized spliced RNA expression matrix and used Wilcoxon rank-sum test to perform DEG analysis. We further used Benjamini & Hochberg method to perform multiple-test correction. Meanwhile, we also calculated each gene expression difference between *Ddx43^KI/KI^* and *Ddx43^+/+^* samples using following equation.

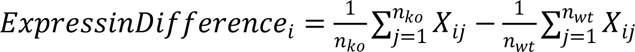

#### Weighted Gene Co-expression Network Analysis (WGCNA)

To prevent the influence of batch effects between *Ddx43^KI/KI^* samples and *Ddx43^+/+^* samples. We only extract the cells of STids from *Ddx43^+/+^* samples and used the downregulated genes in sub-cluster 0 and sub-cluster 1 to perform WGCNA^73^ analysis. Unsigned pearson correlation was first used to calculate pairwise correlations between genes. Next, pairwise topological overlap was calculated with a power of 2 based on a fit to scale-free topology. Co-expression modules comprised of correlated genes with high topological overlap were then identified using the cutreeDynamic function in the dynamicTreeCut R package, with the following parameters: deepSplit = 2, pamRespectsDendro = TRUE, minClusterSize = 50. We applied AUCell to quantify the activity of each module in each cell from *Ddx43^+/+^* and *Ddx43* mutant samples. To visualize the gene expression pattern of modules, we extract the top 20 gene with the highest gene connectivity computed through *intramodularConnectivity* function, and visualize their expression pattern through a heatmap with the cells are ordered according to the velocity time.

### Driver Gene Identification Using Random Walk with Restart algorithm

Guilt-by-association analysis have been widely used to identify possible driver candidates relate to the phenotype of interests. Specifically, we hypothesize that the genes proximal to the four co-expression modules probably could be regarded as the key genes. Therefore, we need to define a topological distance metric to assess the distance between all genes and the four modules respectively, and aggregate distance information to derive a ranking list which represents the global distance of each genes to the four modules. We used Random walk with restart (RWR)^95^, a network diffusion algorithm to measure the distance. Different from conventional random walk methods, RWR introduces a pre-defined restart probability at the initial genes for every iteration, which can take into consideration both local and global topological connectivity pattern within the network to fully exploit the underlying direct or indirect relations between nodes. First, we converted the gene topological overlapped matrix (TOM) derived from WGCNA analysis into gene level transition matrix as follow.

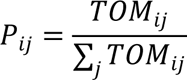

Formally, TOM denote the topological overlapped matrix which represents the adjacency matrix of a gene co-expression network with m genes. The matrix P describes the probability of a transition from gene i to gene j. Next, let 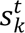 be an m-dimensional distribution vector in which each element stores the probability of each node being visited after t iterations in the random walk process, with the starting points set from the k th module genes. Then RWR from node i can be defined as:

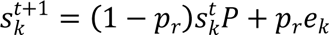

Where *e_k_* stands for an m-dimensional standard basis vector with (*e_k_*)*_i_*(*i* ϵ *module_k_*) = 1 , indicate whether the genes belong to the specific module (e.g., brown module). And *p_r_* stands for the pre-defined restart probability, which actually controls the relative influence between module local and global topological information in the diffusion process. We used following metric to stop the iteration process.

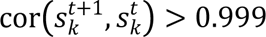

That means once the 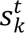 reach a stationary distribution, we stop the iterations. We referred the stationary distribution as the “steady states”, and each element of the steady states vector can be regarded as the inverse of topological distance to the modules. We then aggregated steady states probability vector into the joint association vector defined as follow:

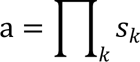

Using the association vector *a*, we could rank all genes and selected the top 30 genes regarded as the proximal genes to all four modules.

## Ethics approval

All experiments involving mice were conducted according to the guidelines of the Institutional Animal Care and Use Committee of Nanjing Medical University.

## Authors’ contributions

K.Z., T.N. and Y.-Q,C. conceived and supervised the project. H.-H.T., W.-X.W. L.Y., Y.-Q, C., T.N., and K.Z. designed the work. H.-H.T., C.-J. Z., Y.-F.W., S.Z., P.-L.Y., and R.G performed experiments. W.-X.W., and W.C. performed single-cell RNA-seq and bioinformatics analyses. H.-H.T., W.-X.W., L.Y., Y.-Q, C., T.N., and K.Z. interpreted the experimental and analytical data. K.Z., H.-H.T., and W.-X.W. wrote the manuscript, and T.N. revised it. All authors discussed and approved the final manuscript.

