## Extended Data for "Single-Cell RNA-seq Uncovers Dynamic Processes Orchestrated by RNA-Binding Protein DDX43 in Chromatin Remodeling during Spermiogenesis"

**Extended Data Fig.S1 to S11**

### Single-Cell RNA-Seq Uncovers Dynamic Processes Orchestrated by RNA-Binding Protein DDX43 in Chromatin Remodeling during Spermiogenesis

Huanhuan Tan[1,†], Weixu Wang[2,†], Chongjin Zhou[1,†], Yanfeng Wang[1], Shu Zhang[1], Pinglan Yang[1], Rui Guo[1], Wei Chen[2], Lan Ye[1], Yiqiang Cui[1,*], Ting Ni[2,*], Ke Zheng[1,*]

[1]State Key Laboratory of Reproductive Medicine, Nanjing Medical University, Nanjing 211166, China; [2]State Key Laboratory of Genetic Engineering, Collaborative Innovation Center of Genetics and Development, Human Phenome Institute, Shanghai Engineering Research Center of Industrial Microorganisms, School of Life Sciences and Huashan Hospital, Fudan University, Shanghai 200438, China

† These authors contributed equally to this work

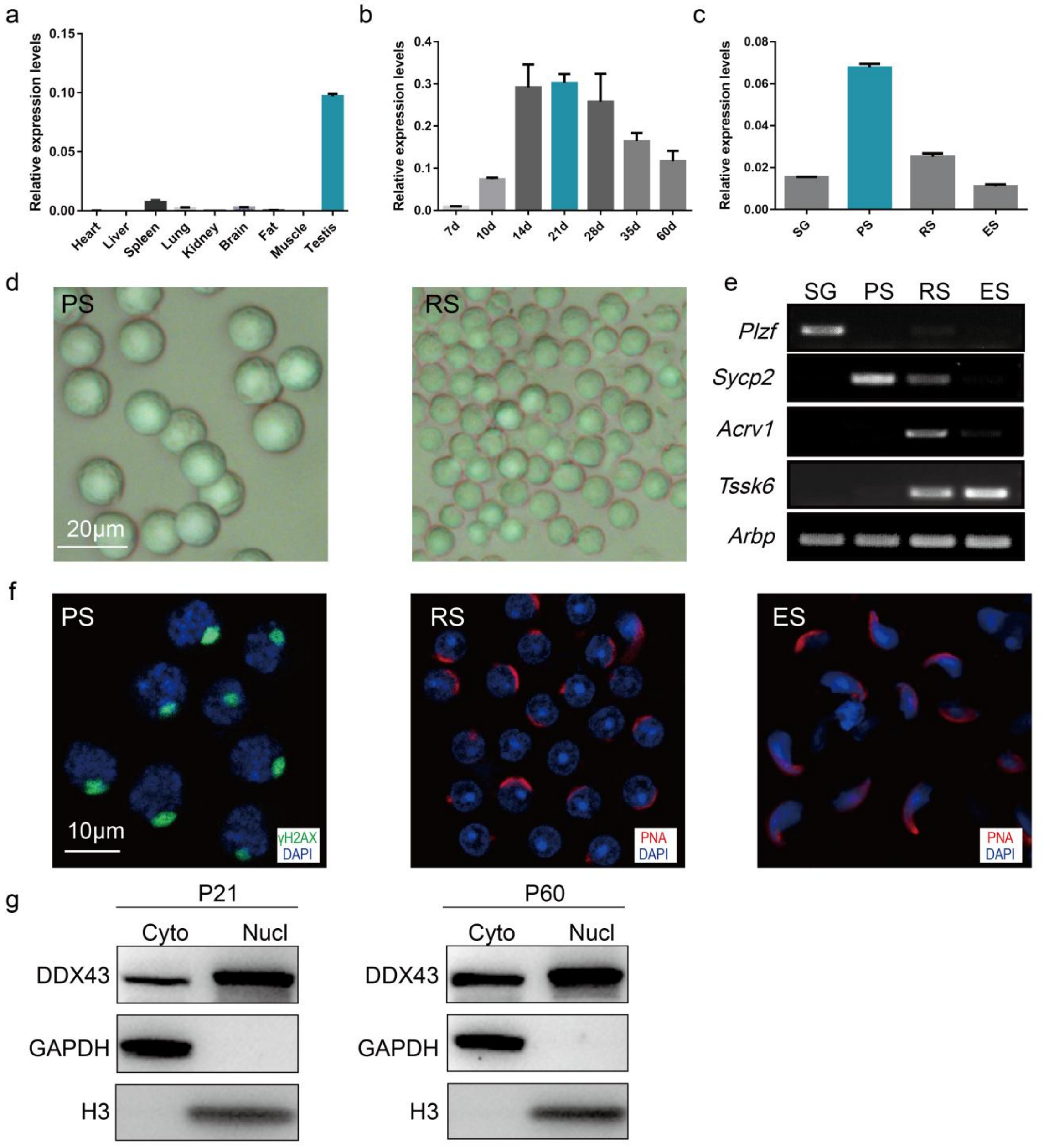


**Extended Data figure 1. *Ddx43* expression and purified germline populations. Related to Figure 1.**

(a-c) Quantitative RT-PCR (qRT-PCR) analyses of *Ddx43* mRNA transcripts from multiple adult mouse tissues (a), testis tissues collected from mice at different ages indicated (b), and various types of isolated spermatogenic cell populations (c). SG, spermatogonia (approximately 85% purity); PS, pachytene spermatocytes (approximately 90% purity); RS, round spermatids (approximately 90% purity); ES, elongating spermatids (approximately 85% purity). Results are normalized to *Rplp0*

(*36b4*). Data presented are mean ± s.d. from three independent experiments.

(d) Morphology analyses of purified pachytene spermatocytes (PS) and round spermatids (RS). Scale bar is indicated.

(e) RT-PCR analyses of key marker genes further confirm the identity of cell populations, including *Plzf* for spermatogonia, *Sycp2* for meiotic spermatocytes, *Acrv1* for round spermatids, *Tssk6* for post-meiotic spermatids. *Arbp* serves as a control.

(f) Immunofluorescence staining of specific marker protein γ-H2AX (green) and PNA (red) for isolated pachytene spermatocytes (PS), round spermatids (RS) and elongating spermatids (ES). DNA was counterstained with DAPI. Scale bar is indicated.

(g) Protein levels of DDX43 in cytoplasmic and nuclear fractions of mouse testes at ages indicated. GAPDH and histone H3 are cytoplasmic and nuclear positive markers, respectively.

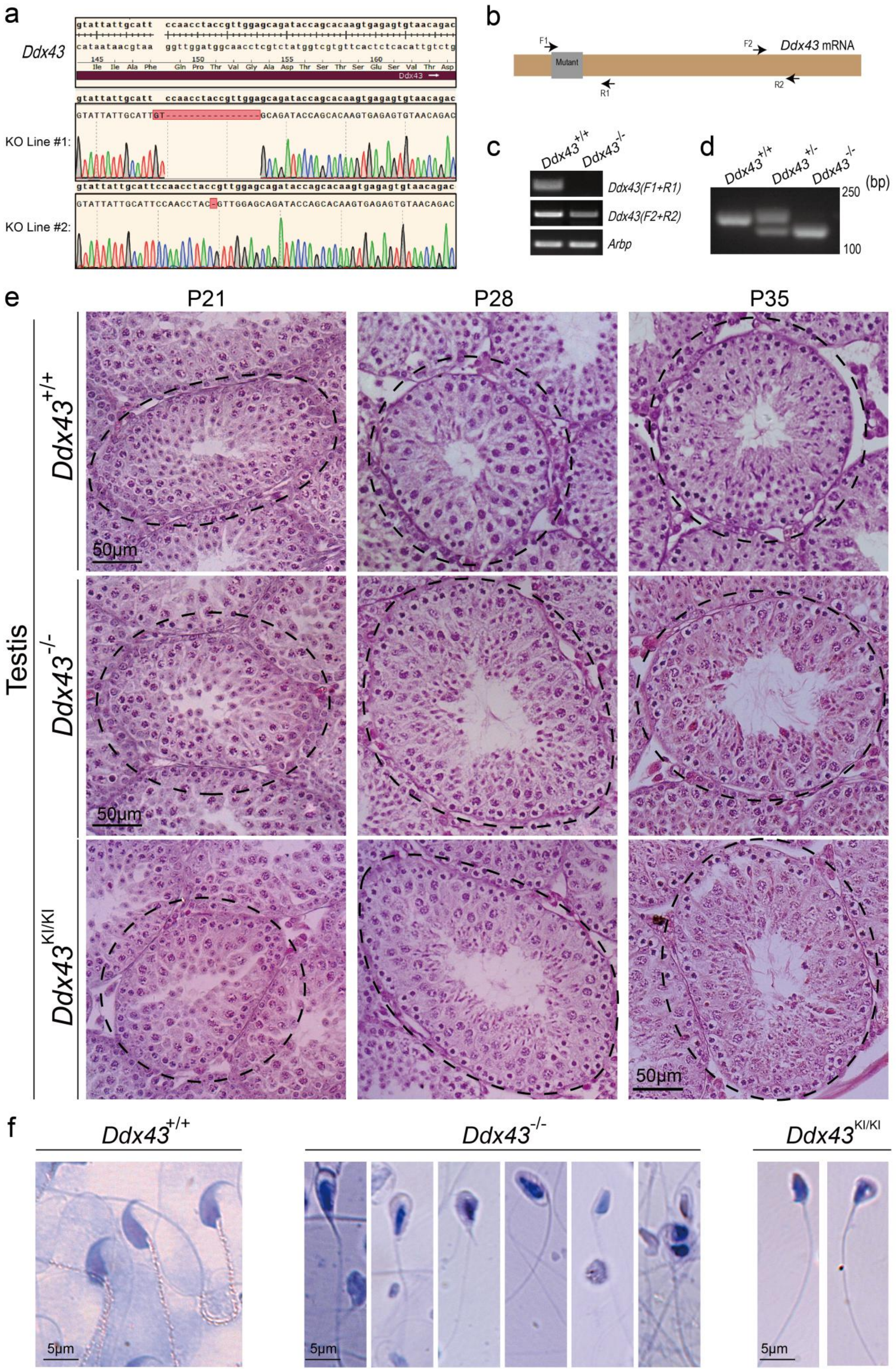
a
Ddx43
KO Line #1:
KO Line #2:
b
Ddx43 mRNA
Mutant
F1
R1
F2
R2
c
Ddx43+/+
Ddx43-/-
Ddx43(F1+R1)
Ddx43(F2+R2)
Arbp
d
Ddx43+/+
Ddx43+/-
Ddx43-/-
250
100
(bp)
e
P21
P28
P35
Testis
Ddx43+/+
Ddx43-/-
Ddx43KI/KI
50μm
f
Ddx43+/+
Ddx43-/-
Ddx43KI/KI
5μm

**Extended Data figure 2. Sequence validation and phenotypic analyses in *Ddx43* mutant mice. Related to Figure 2.**

(a) Chromatogram of Sanger sequencing in *Ddx43*$^{+/+}$ and *Ddx43*$^{–/–}$ mice, showing the successful generation of *Ddx43* mutant.

(b) Primer design of RT-PCR for identifying wild-type and mutant *Ddx43* transcripts. Primers F1/R1 and F2/R2 are positioned relative to sgRNA-1 and -2's targeting sites.

(c) Based on the RT-PCR and primer pairs shown in panel B, full-length transcripts are present in wild-type testes, but not in those of *Ddx43*$^{–/–}$ mice, which only contain truncated *Ddx43* mRNA. *Arbp* serves as a control.

(d) Genotyping PCR products of *Ddx43*$^{+/+}$, *Ddx43*$^{+/-}$ and *Ddx43*$^{–/–}$ mice, respectively. These products were confirmed by Sanger sequencing, shown identical in panel A.

(e) Hematoxylin and Eosin (H&E) staining of testes sections from *Ddx43*$^{+/+}$, *Ddx43*$^{KI/KI}$ and *Ddx43*$^{–/–}$ mice at indicated postnatal time points. Note the germ cell defects emerged in P28, when the nuclei of spermatids become flattened and start to elongate along with condensation of chromatin. Scale bar is indicated.

(f) Acidic Aniline staining of epididymal sperm from adult mice. In contrast to the canonical hook-shaped appearance of sperm heads in *Ddx43*$^{+/+}$ mice, all sperm heads in *Ddx43*$^{–/–}$ and *Ddx43*$^{KI/KI}$ mice are amorphous with a smaller and more rounded shape. Scale bar is indicated.

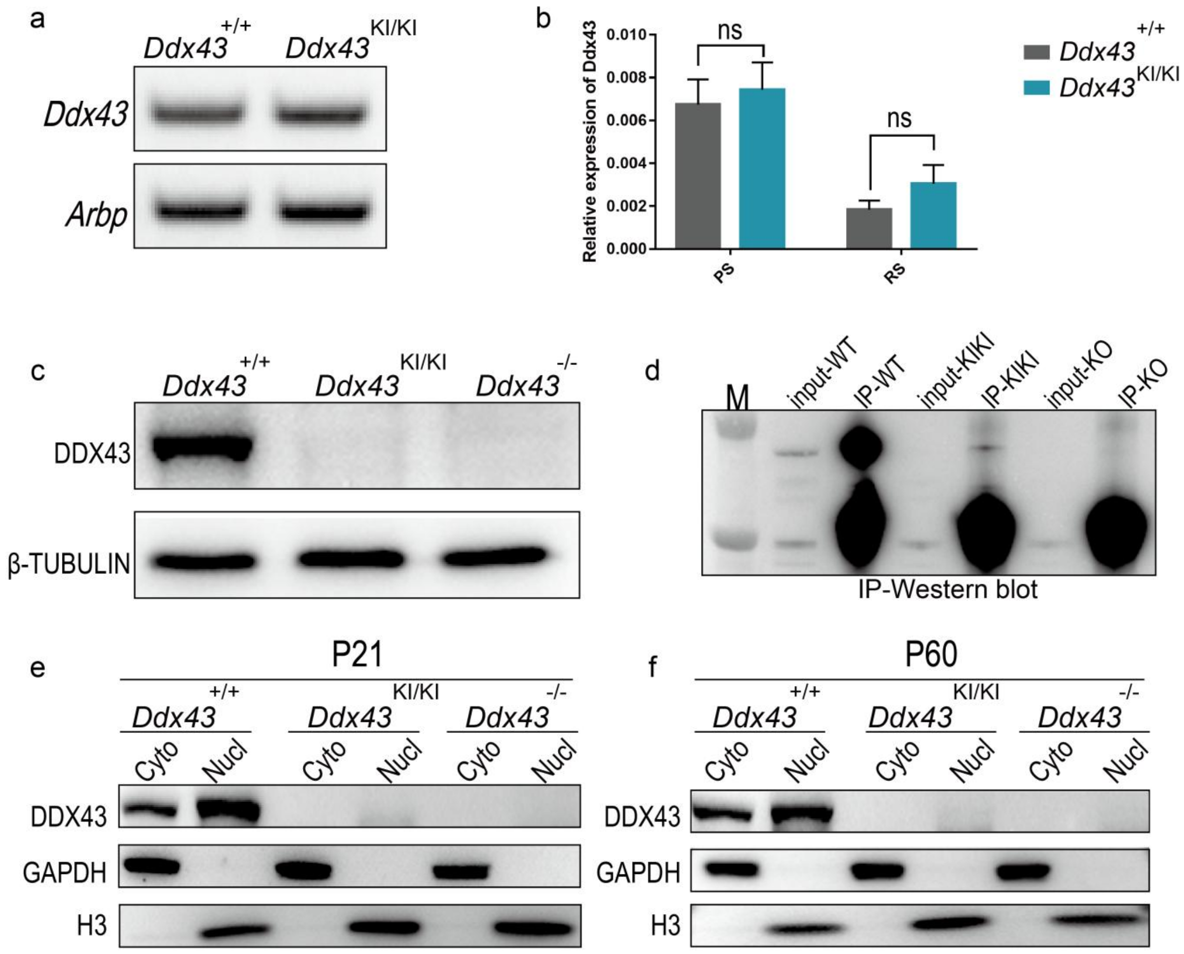


**Extended Data figure 3. DDX43 expression in *Ddx43*$^{KI/KI}$ testes.**

(a) RT-PCR assessment of *Ddx43* mRNA transcripts from adult *Ddx43*$^{+/+}$ and *Ddx43*$^{KI/KI}$ testes. *Arbp* serves as a control.

(b) qRT-PCR assessment of *Ddx43* mRNA transcripts in germ cells isolated from adult *Ddx43*$^{+/+}$ and *Ddx43*$^{KI/KI}$ testes. PS, pachytene spermatocytes; RS, round spermatids. Results are normalized to *Rplp0* (*36b4*). Data presented are mean ± s.d. from three independent experiments.

(c) Western blot analyses of DDX43 protein in lysates from adult testes of indicated genotypes. β-TUBULIN serves as an internal loading control.

(d) Western blot analyses of DDX43 protein in its immunoprecipitation (IP) lysates from testes of indicated genotypes. The *Ddx43*$^{KI/KI}$ mutant protein is detectable at a greatly reduced level after IP enrichment.

(e, f) Western blot analyses of DDX43 protein in cytoplasmic and nuclear fractions from P21 and P60 testes of indicated genotypes. GAPDH and histone H3 serve as markers for cytoplasm and nucleus, respectively.

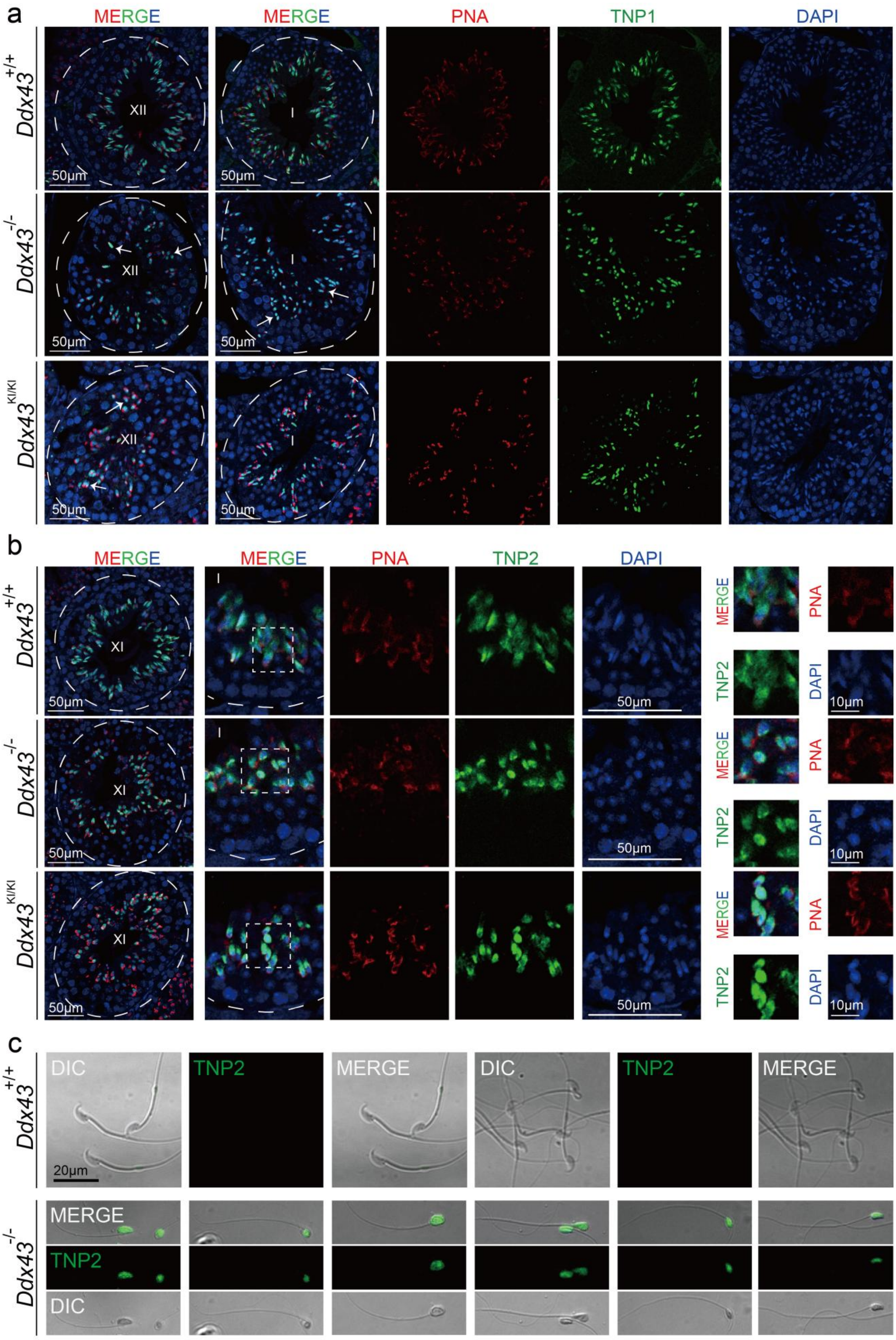
a
MERGE
MERGE
PNA
TNP1
DAPI
Ddx43+/+
Ddx43-/-
Ddx43KI/KI
XII
I
50μm
b
MERGE
MERGE
PNA
TNP2
DAPI
XI
I
50μm
10μm
c
DIC
TNP2
MERGE
20μm
Ddx43+/+
Ddx43-/-

**Extended Data figure 4. Localization of TPs in adult testes. Related to Figure 3.**

(a) Co-immunofluorescence staining of TNP1 (green) and PNA (red) in testes sections from adult mice of indicated genotypes at stages XII-I, showing comparable expression abundance between *Ddx43*$^{+/+}$, *Ddx43*$^{KI/KI}$ and *Ddx43*$^{-/-}$ mice. The elongating spermatids nuclei in *Ddx43*$^{+/+}$ mice were canonical hook-shaped and thinner along with differentiation; yet, the nuclei of spermatids were deformed as rod-shaped or round-like in *Ddx43*$^{KI/KI}$ and *Ddx43*$^{-/-}$ mice. White arrows mark abnormal spermatids. DNA was counterstained with DAPI. Scale bar is indicated.

(b) Co-immunostaining of TNP2 (green) and PNA (red) in *Ddx43*$^{KI/KI}$ and *Ddx43*$^{-/-}$ mice revealed anomalous morphological structures similar to panel A. DNA was counterstained with DAPI. Scale bar is indicated.

(c) Representative immunostaining images of TNP2(green) and PNA (red) in sperm from indicated genotypes. Note the abnormal head shape and TNP2 localization, indicative of less condensed mutant sperm. Scale bar is indicated.

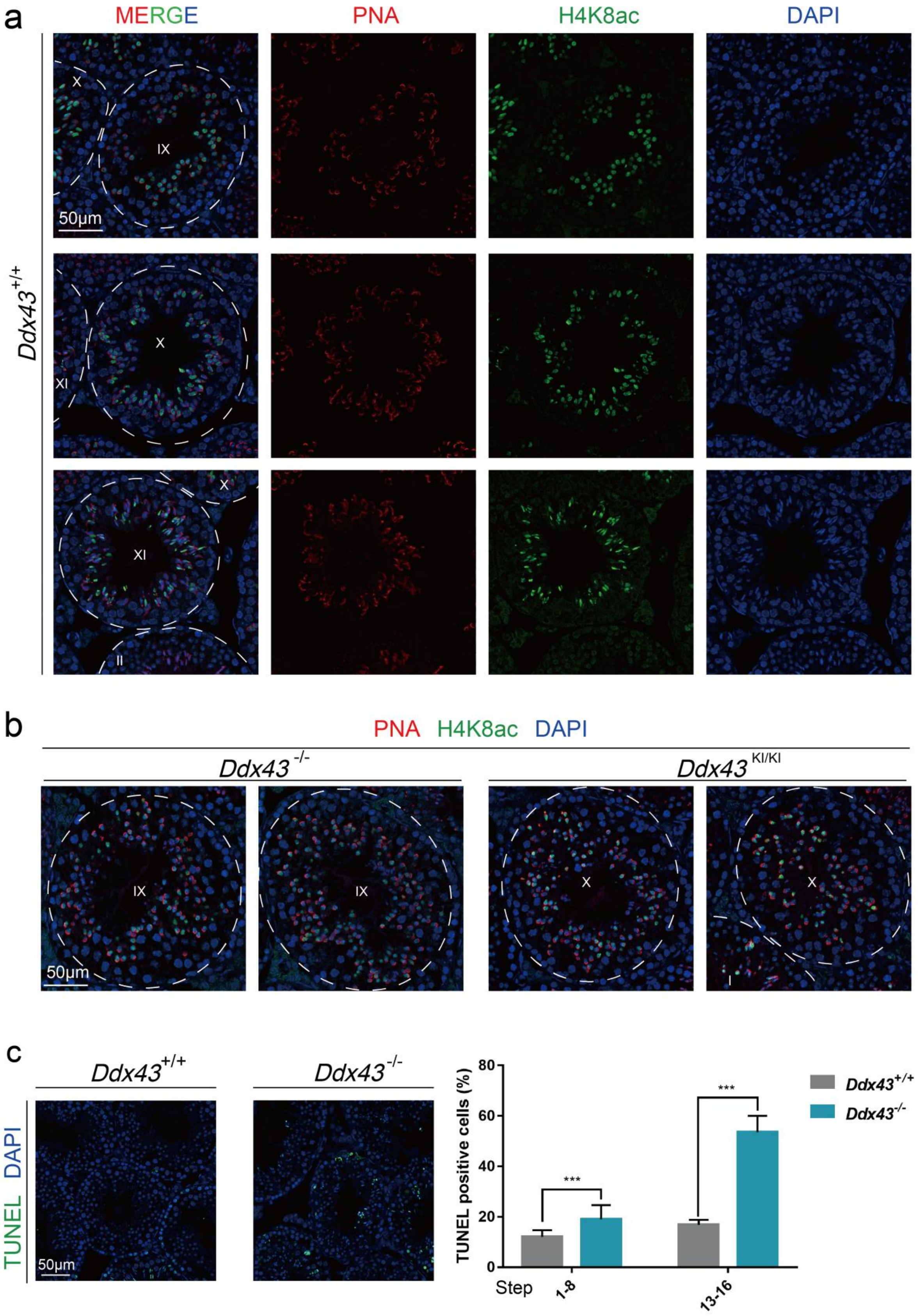


**Extended Data figure 5. Localization of H4K8ac in adult testes. Related to Figure 3.**

(a, b) Immunofluorescence analyses of H4K8ac in adult *Ddx43*$^{+/+}$, *Ddx43*$^{KI/KI}$ and *Ddx43*$^{-/-}$ seminiferous tubules. In *Ddx43*$^{+/+}$ sections, H4K8ac emerges in step 9 spermatids at stage IX, peaks in step 10-11 spermatids at stage X-XI, disappears from step 12 onward. And the comparable expression intensity in mutant spermatids indicates DDX43 protein is dispensable for H4 acetylation during spermiogenesis. Stage numbers and scale bars are indicated.

(c) Representative images (left) and statistics of positive cells (right) in TUNEL assay on adult *Ddx43*$^{+/+}$ and *Ddx43*$^{-/-}$ sections. Note the increased signals from the mutant elongating spermatids, indicative of increased cell death. Data are presented as mean±s.d. from three independent experiments. Scale bar is indicated.

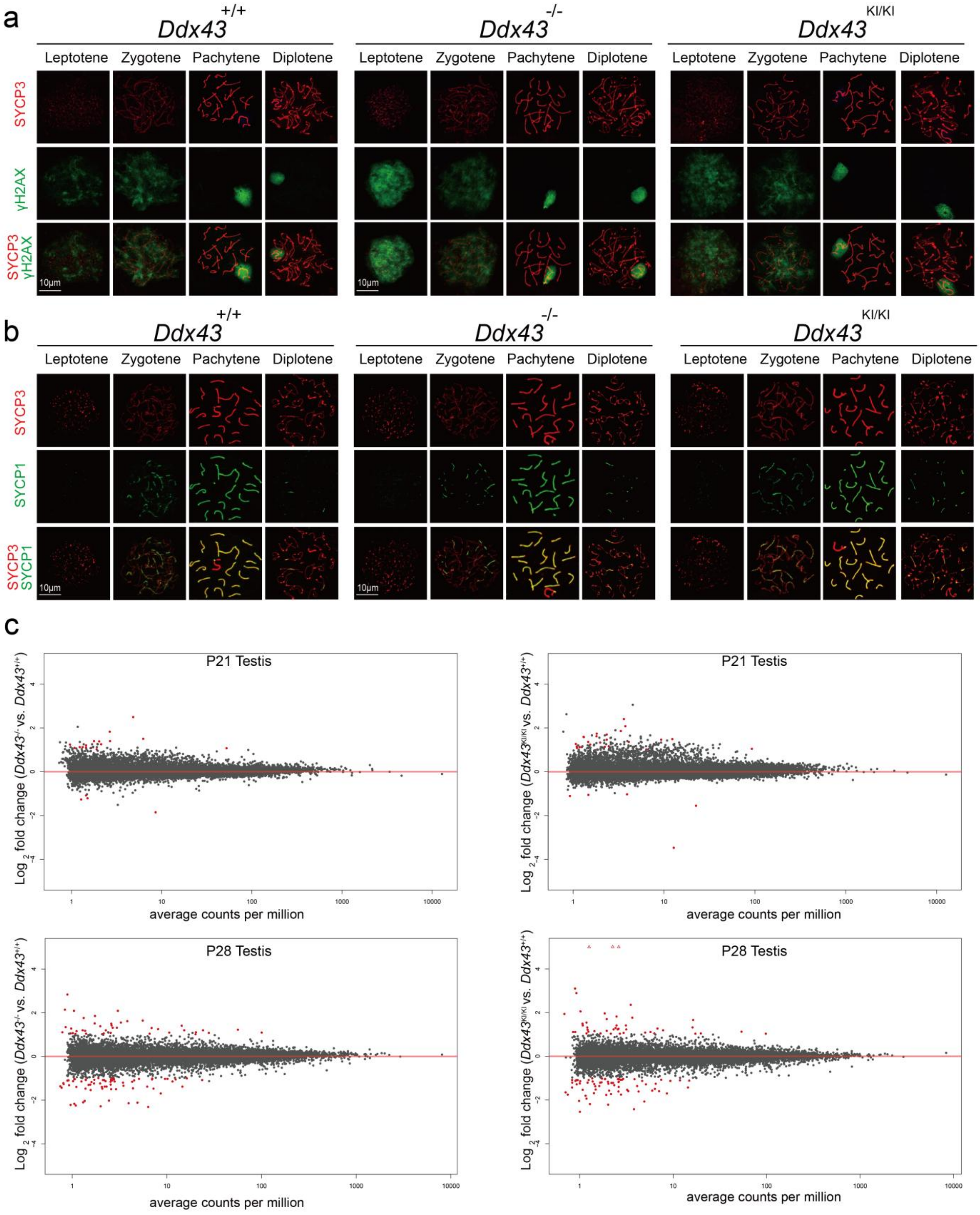


**Extended Data figure 6. Analyses of spermatocyte chromosome spread and testicular transcriptome.**

(a, b) Representative images of co-immunostaining of spermatocyte spread nuclei from *Ddx43*$^{+/+}$, *Ddx43*$^{KI/KI}$ and *Ddx43*$^{–/–}$ adult testes across four different stages (leptotene, zygotene, pachytene and diplotene) of meiotic prophase I. (a) γH2AX (green) and

SYCP3 (red) show dynamic patterns of DNA double strand breaks (DDSBs). The formation of DSBs in meiosis can be visualized by γH2AX, which emerges and spreads over the leptotene spermatocytes, persists into zygotene stage, reduced in early pachytene stage, and is confined to the XY body in mid-late pachynema and diplonema.
(b) Co-immunostaining of SYCP1 (green) and SYCP3 (red) shows the developmental process of homologous chromosome synapsis, which initiates at the zygotene stage and completes at the onset of the pachytene stage. Thus, all autosome regions can be immunolabeled by SYCP1 and SYCP3, which are otherwise confined within the short pseudoautosomal region of the sex chromosomes. DNA was counterstained with DAPI. Scale bar is indicated.
(c) Scatter plot of differentially expressed transcripts in *Ddx43* mutant testes at P21 (left) and P28 (right) compared with age-matched *Ddx43*$^{+/+}$ testes. Genes showing altered expression with $p < 0.05$ and > 1.5 fold changes are colored red.

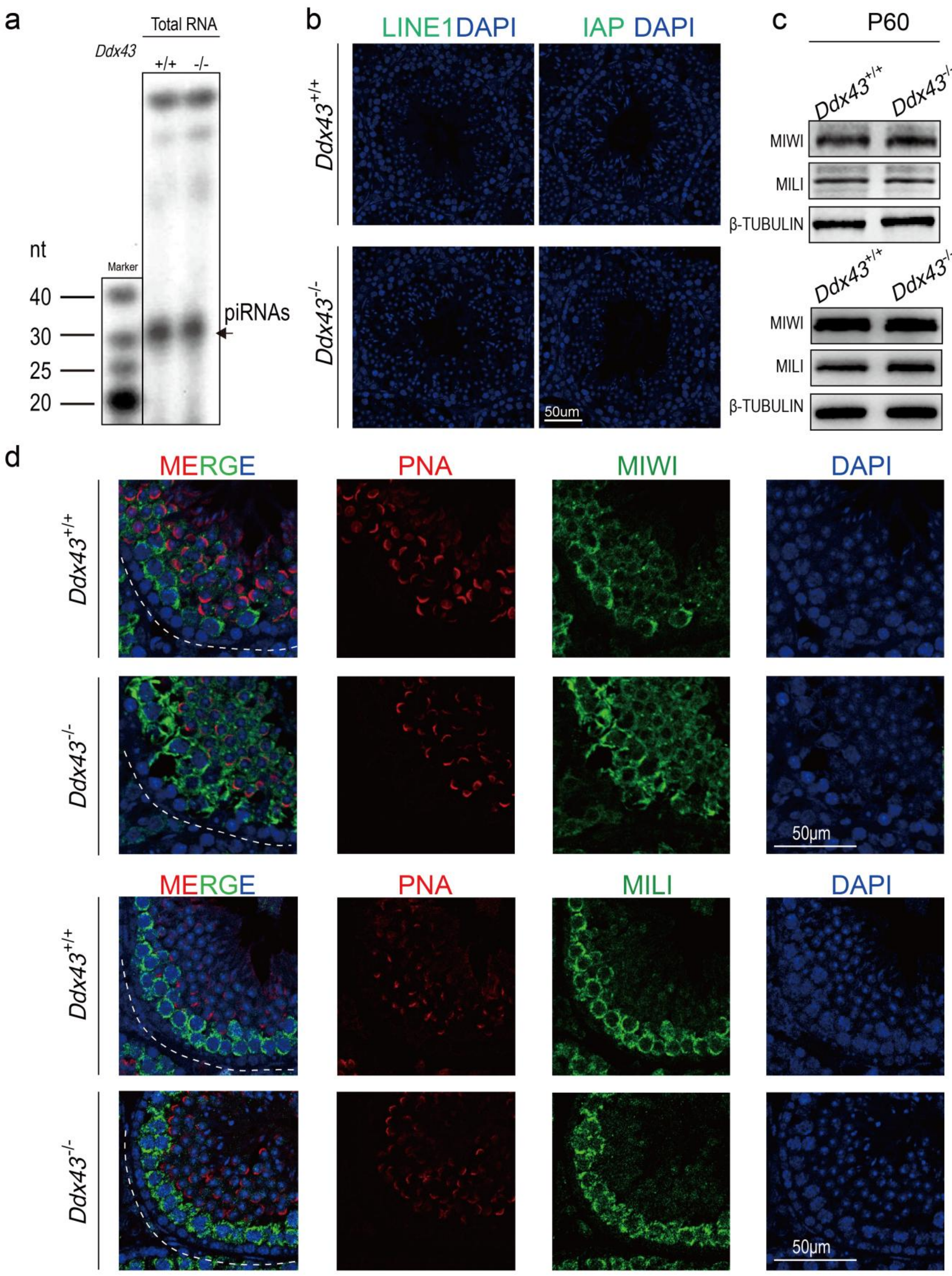


**Extended Data figure 7. Inspection of the piRNA pathway in testes.**

(a) Pachytene piRNA abundance is not reduced in mutant testes from adult *Ddx43*$^{-/-}$ mice. $^{32}$P-end-labelled total RNAs are separated for autoradiography by denaturing polyacrylamide gel electrophoresis.

(b) Immunofluorescence detection of LINE1 ORF1p and IAP from *Ddx43*$^{+/+}$ and *Ddx43*$^{-/-}$ P60 testes. DNA was counterstained with DAPI. Scale bar is indicated.

(c) Western blot analyses of MIWI and MILI in lysates from adult *Ddx43*$^{+/+}$ and *Ddx43*$^{-/-}$testes. β-TUBULIN serves as an internal loading control.

(d) Immunofluorescence analyses of piRNA pathway components MIWI (green) and MILI (green) in *Ddx43*$^{+/+}$ and *Ddx43*$^{-/-}$ testes. DNA was counterstained with DAPI. Scale bar is indicated.

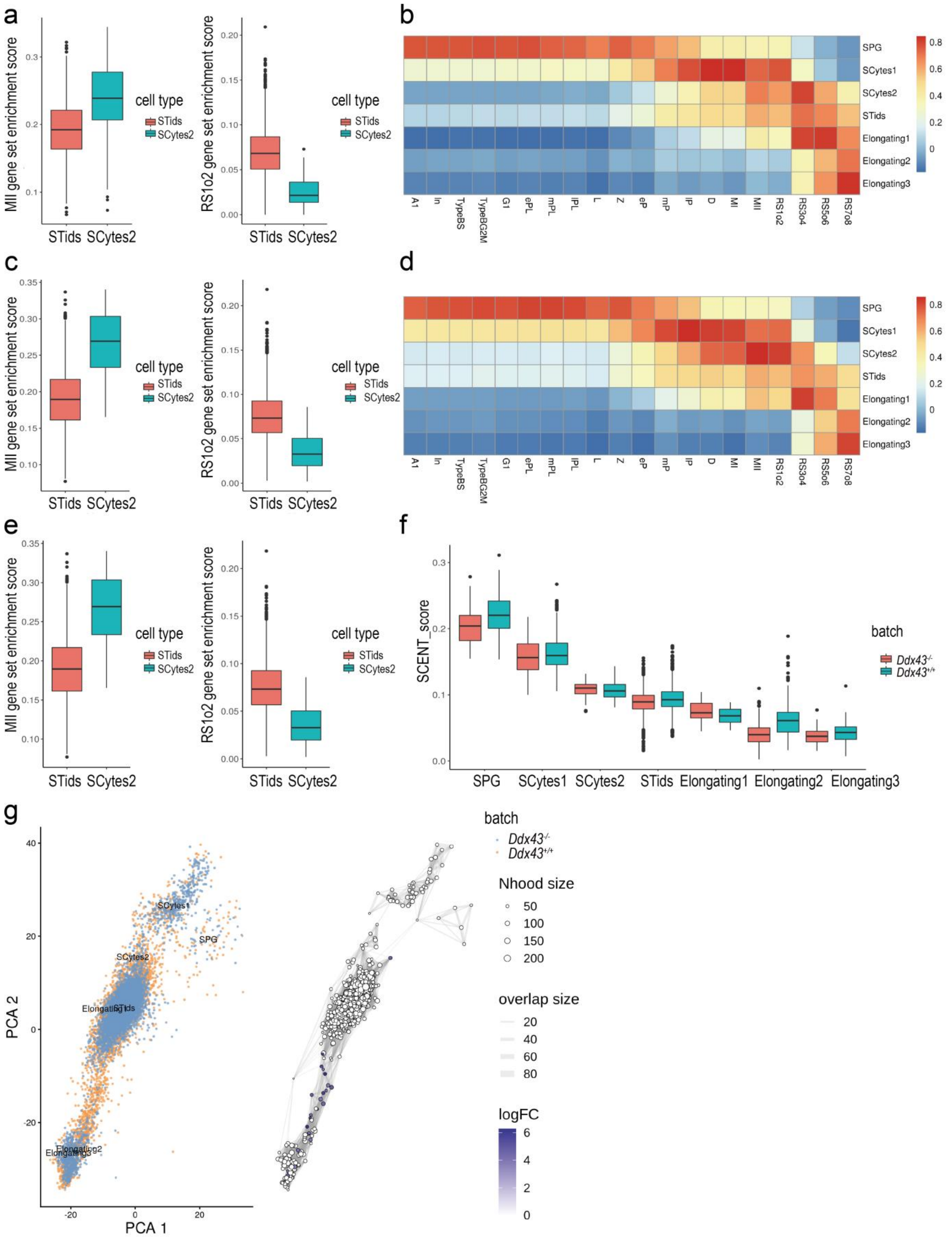


**Extended Data figure 8. Other characteristic analysis of Single Cell RNA-Seq in *Ddx43*$^{+/+}$ and *Ddx43* mutant testes. Related to Figure 4.**

(a, c, e) Box plot of metagene expression in SCytes2 and STids in *Ddx43*$^{+/+}$(a), *Ddx43*$^{KI/KI}$ (c) and *Ddx43*$^{-/-}$ (e) . The gene sets are curated from previous differential expression study between MII and Spermatids stage1 to stage 2[1]. The metagene expression values are determined through AUCell[2].

(b, d) The heatmap of our annotated *Ddx43*$^{KI/KI}$ (b) and *Ddx43*$^{-/-}$ (d) single-cell expression profile and the published flow-sorted single cell gene expression profile in cell type gene expression spearman correlation. Gene ontology analysis of the four subtypes from STids.

(g) A neighborhood graph of the results from Milo differential abundance testing (right panel). Nodes are neighborhoods, colored by their log fold change across genotypes. Non-differential abundance neighborhoods (P-value > 0.05) are colored white, and sizes correspond to the number of cells in each neighborhood. Graph edges depict the number of cells shared between neighborhoods. The layout of nodes is determined by the position of the neighborhood index cell in the PCA (left panel).

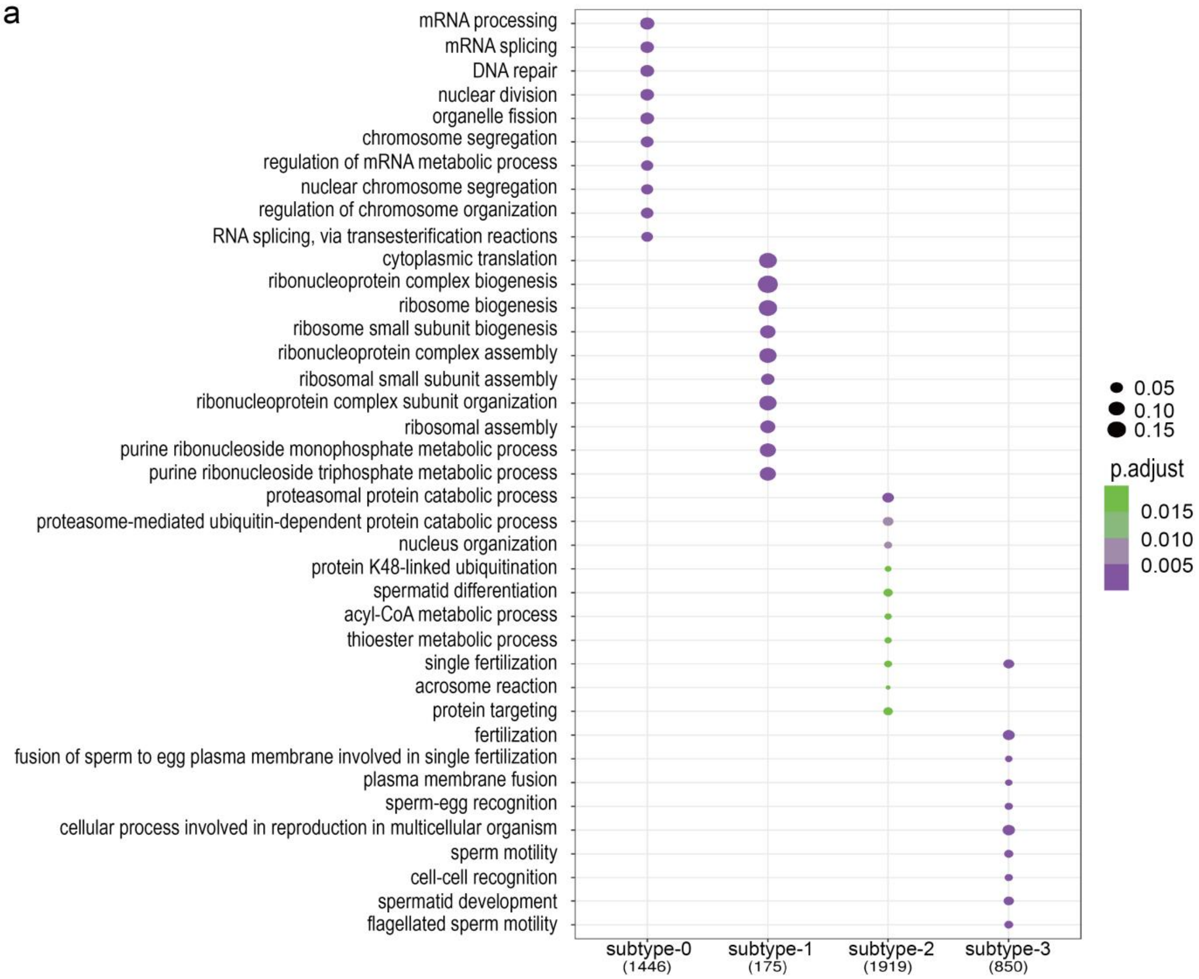


**Extended Data figure 9. Characterize gene expression pattern of four cellular states in STids. Related to Figure 5.**

(a) Go enrichment of the marker genes of four cellular states of STids.

a

eCLIP sequencing library processing summary metrics

| Sample | | Raw reads | short reads | pcr_dup | clean reads | Total clean reads | Total mapped | Total Uniquely mapped | Total Multiple mapped |
|---|---|---|---|---|---|---|---|---|---|
| SMInput1 | 50999456 | 1684654 (3.30%) | 1684654 (3.30%) | 11220851 (22.00%) | 38093951 (74.69%) | 75818467 | 61133433 (80.63%) | 40448651 (66.16%) | 20684782 (33.84%) |
| | 50638651 | 2410635 (4.76%) | 2410635 (4.76%) | 10503500 (20.74%) | 37724516 (74.50%) | | | | |
| DDX43 IP1 | 56004349 | 2746478 (4.90%) | 2746478 (4.90%) | 13709401 (24.48%) | 39548470 (70.62%) | 75452592 | 50062926 (66.35%) | 41894388 (83.68%) | 8168538 (16.32%) |
| | 55210178 | 4103965 (7.43%) | 4103965 (7.43%) | 12659743 (22.93%) | 38446470 (69.64%) | | | | |
| SMInput2 | 50868017 | 1706389 (3.35%) | 1706389 (3.35%) | 10584828 (20.81%) | 38576800 (75.84%) | 77994940 | 61447833 (78.78%) | 40243788 (65.49%) | 21204045 (34.51%) |
| | 50770778 | 4029196 (7.94%) | 4029196 (7.94%) | 9865790 (19.43%) | 36875792 (72.63%) | | | | |
| DDX43 IP2 | 56964723 | 1504015 (2.64%) | 1504015 (2.64%) | 18107359 (31.79%) | 37353349 (65.57%) | 69684187 | 48439019 (69.51%) | 39981788 (82.54%) | 8457231 (17.46%) |
| | 56825817 | 3833629 (6.75%) | 3833629 (6.75%) | 11981027 (21.08%) | 41011161 (72.17%) | | | | |

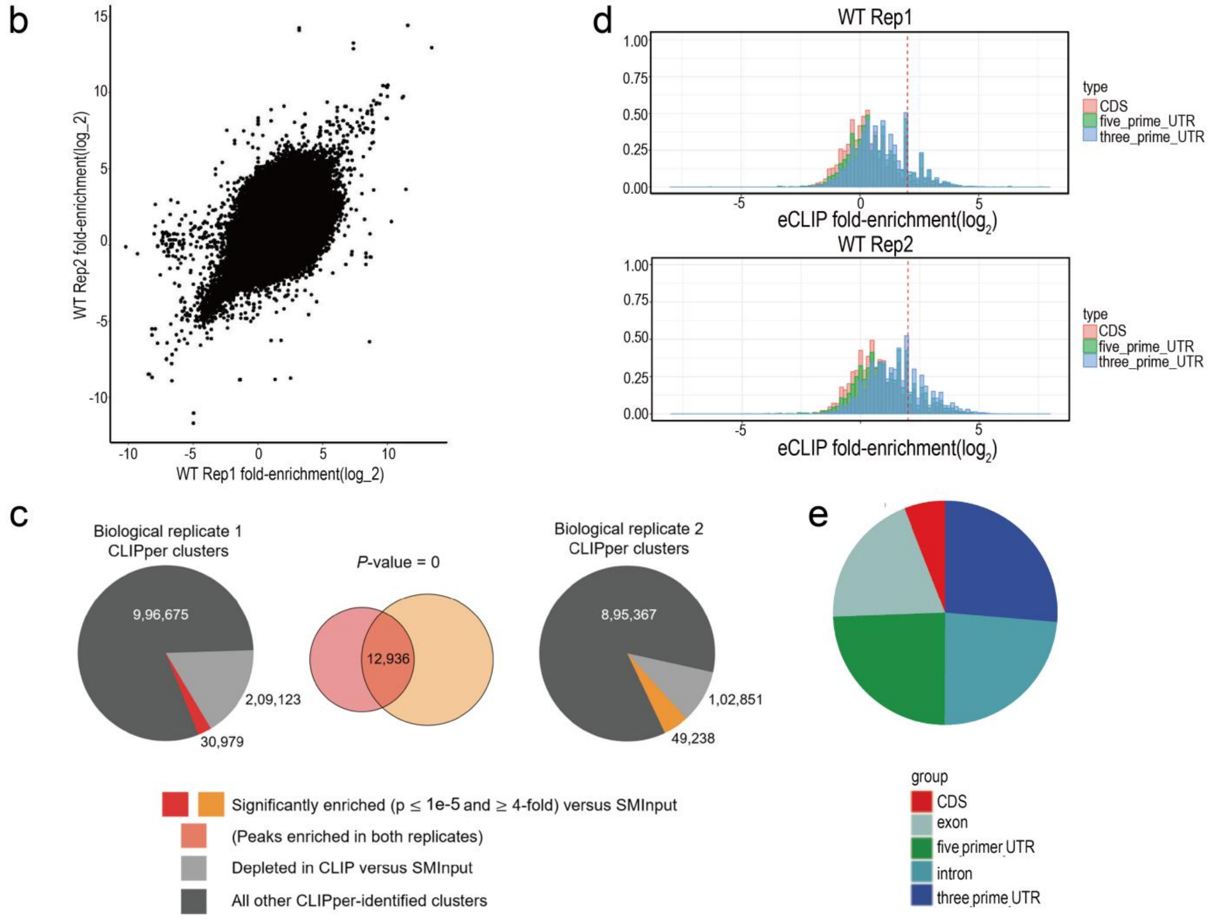

**Extended Data figure 10. Genome-wide mapping of DDX43 eCLIP-seq. Related to Figure 4.**

(a) A table related to eCLIP-seq library processing summary metrics.

(b) Scatter plot indicates correlation between DDX43 biological replicates based on the fold enrichment.

(c) For each biological replicate, CLIPper (gray) identifies clusters of enriched read density within DDX43 eCLIPs. Based on cluster read density comparisons between eCLIP and paired SMInput, we identified subsets of clusters enriched above SMInput (red/orange), which display high overlap between replicates (center).

(d) Histogram of region-based fold enrichment for DDX43 with paired SMInput as comparison.

(e) Pie chart showing the distribution of significantly enriched replicate peaks for DDX43 eCLIP-seq data.

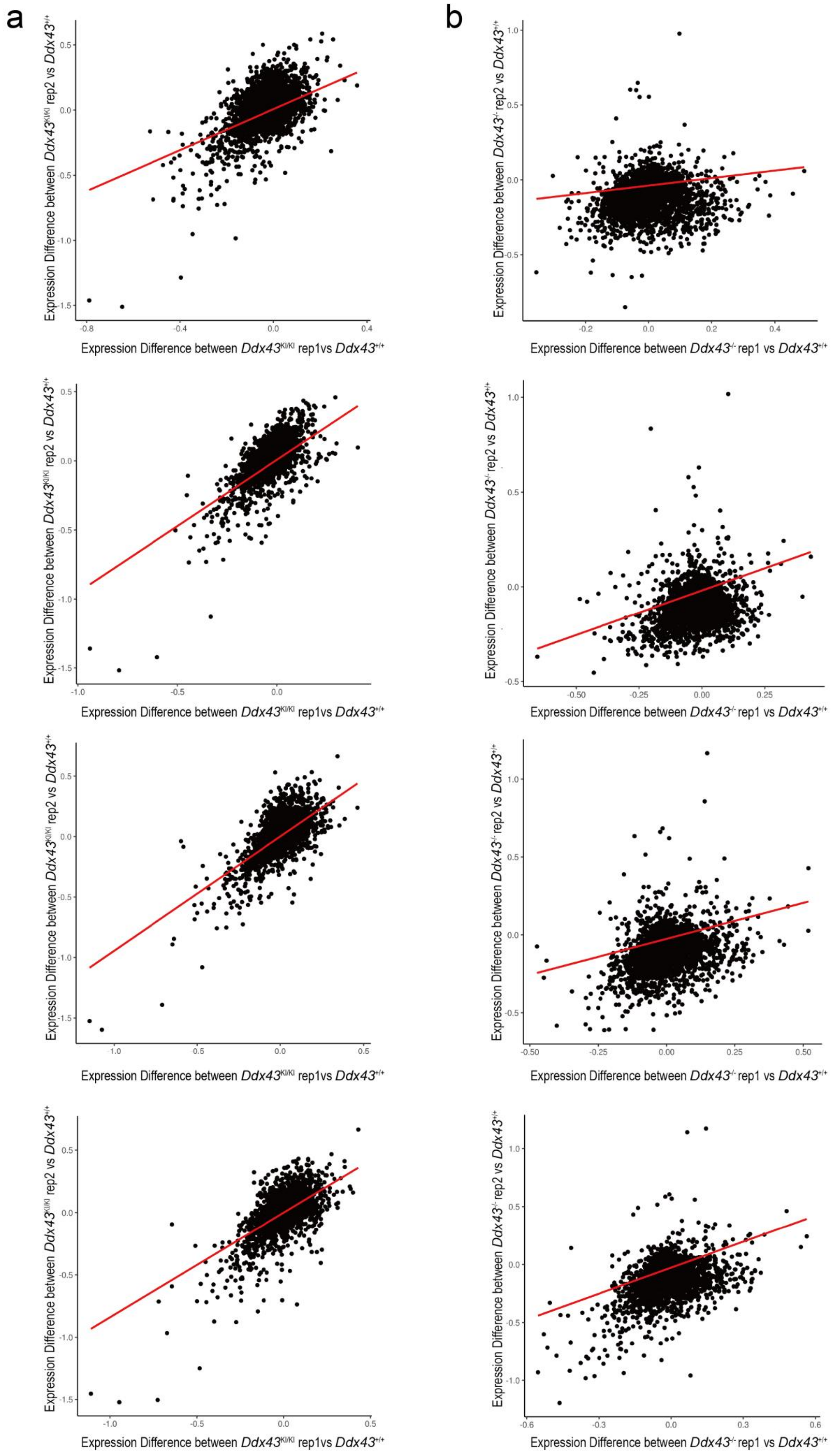
a
Expression Difference between $Ddx43^{KI/KI}$ rep2 vs $Ddx43^{+/+}$
Expression Difference between $Ddx43^{KI/KI}$ rep1vs $Ddx43^{+/+}$
b
Expression Difference between $Ddx43^{-/-}$ rep2 vs $Ddx43^{+/+}$
Expression Difference between $Ddx43^{-/-}$ rep1 vs $Ddx43^{+/+}$

**Extended Data figure 11. Validation of DEG analysis between *Ddx43*$^{+/+}$ and *Ddx43* mutant.**

(a, b) Scatter plot of the Fold Change values of all genes in four sub-clusters. For each *Ddx43* mutant genotype we have two replicates, for each replicate we perform DEG analysis with the wild-type in four sub-clusters respectively, and calculate the fold change values, then comparing whether the Fold Change values are consistence between two replicates.

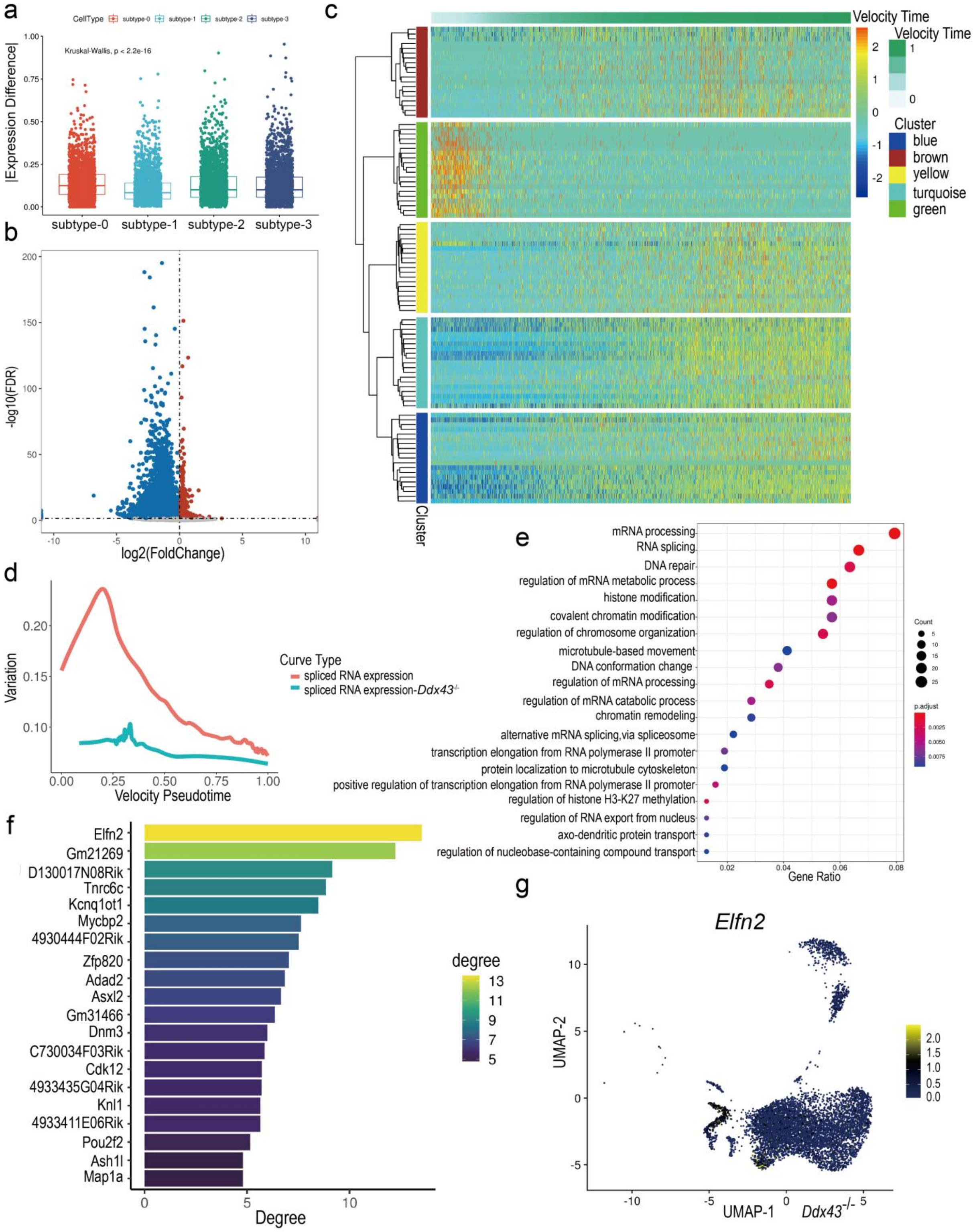


**Extended Data figure 12. Dynamic network analyses using *Ddx43*$^{-/-}$ and wild-type samples**

(a) The boxplot for absolute expression difference value of DDX43 binding targets (characterized by eCLIP-seq) in four cellular states of STids.

(b) Volcano plot showing gene differential expression (*Ddx43*$^{+/+}$ vs *Ddx43*$^{-/-}$) of subtype1.

(c) WGCNA clustering of genes exhibiting down-regulated expression in *Ddx43*$^{-/-}$ samples during subtype-1. Each row represents a gene, and each column represents a single cell, with columns/cells placed in velocity pseudotime order and depicted by a thick colored line (top). Gene expression levels utilize a Z score transformation.

(d) Line plot of green module activity along with velocity pseudotime, module activity is calculated using AUCell.

(e) GO analysis of the genes in green module.

(f) Barplot of the network connectivity of driver candidates. Each bar is colored according to each gene connectivity in the network, which reflects the gene's importance to the biological systems.

(g) Expression patterns of *Elfn2* in *Ddx43*$^{-/-}$ mutant.

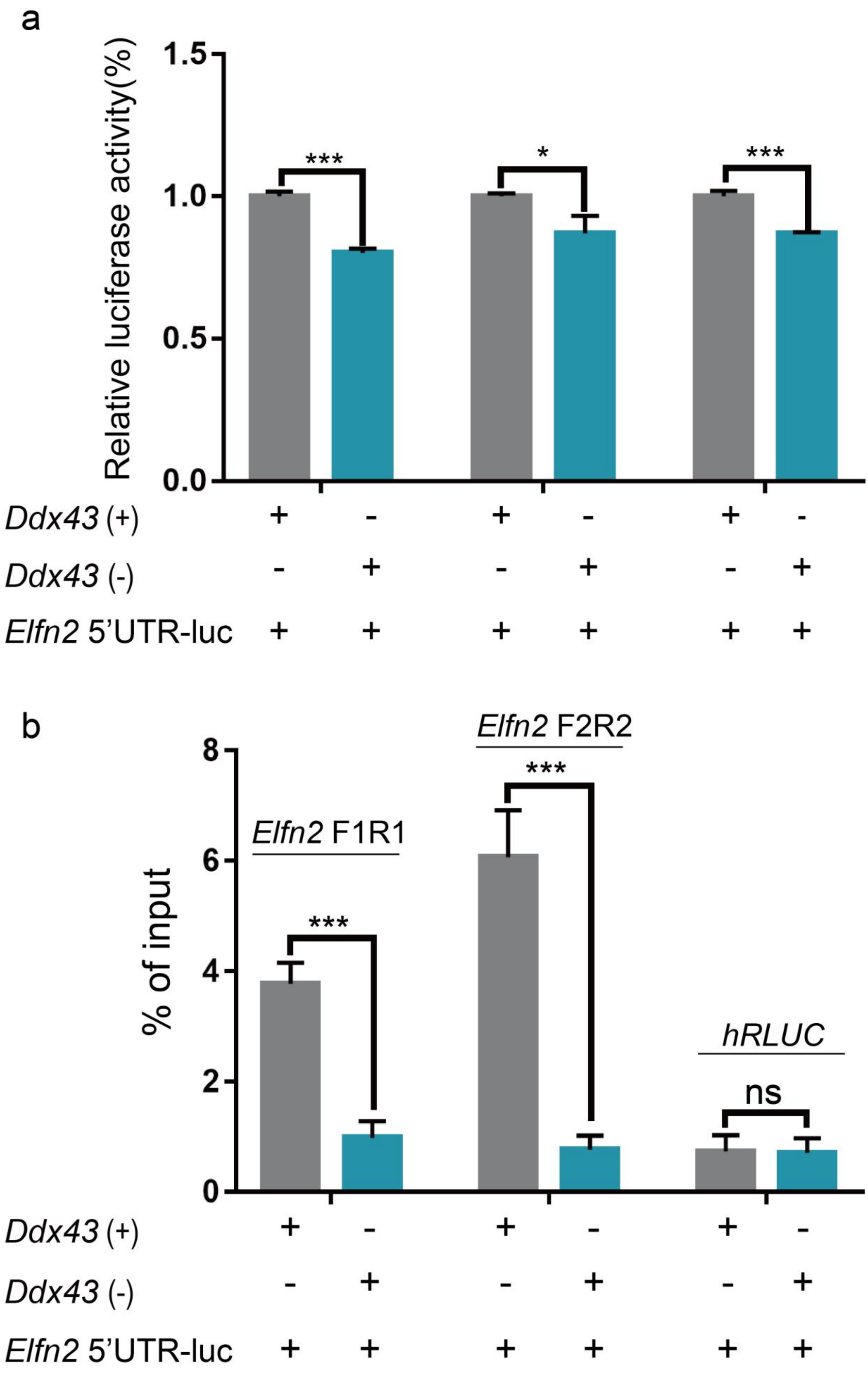
a
Relative luciferase activity(%)
1.5
1.0
0.5
0.0
***
*
***
Ddx43 (+) + - + - + -
Ddx43 (-) - + - + - +
Elfn2 5'UTR-luc + + + + + +
b
% of input
8
6
4
2
0
Elfn2 F1R1
Elfn2 F2R2
hRLUC
***
***
ns
Ddx43 (+) + - + - + -
Ddx43 (-) - + - + - +
Elfn2 5'UTR-luc + + + + + +

**Extended Data figure 13. The validation of *Elfn2* bound by DDX43 in HEK293 cells.**

(a) Dual luciferase reporter assay showing the reporter gene fused with *Elfn2* 5' UTR is regulated by DDX43. **P < 0.01, N = 3.

(b) RNA immunoprecipitation (RIP) was performed using anti-DDX43 antibodies in HEK293 cells transduced as panel F. qRT-PCR detected significant enrichment of DDX43-bound *Elfn2* transcripts in DDX43 immunoprecipitated complexes. Percentage of input is used to calculate binding. ***P < 0.001, N = 3, ns, not significant (Student' s t-test).
